# Chinmo defines the region-specific oncogenic competence in the *Drosophila* central nervous system

**DOI:** 10.1101/2025.05.20.655209

**Authors:** Phuong-Khanh Nguyen, Francesca Froldi, Owen Marshall, Tony D Southall, Louise Y Cheng

**Affiliations:** Sir Peter MacCallum Department of Oncology, The University of Melbourne, VIC 3010, Australia; Peter MacCallum Cancer Centre, Melbourne, VIC 3000, Australia; Department of Anatomy and Physiology, The University of Melbourne, VIC 3010, Australia; Menzies Institute for Medical Research Medical Science Precinct, Hobart TAS, Australia; Department of Life Sciences, Imperial College London, London, UK

**Keywords:** Dedifferentiation, *Drosophila*, malignancy, terminal differentiation

## Abstract

While genetic mutations can promote hyperplastic growth, they do not always result in oncogenic outcomes. We and others have previously identified the transcription factors Nerfin-1 and Lola as inhibitors of dedifferentiation. Here, we investigate how the oncogenic potential of dedifferentiation varies across different neural lineages in the *Drosophila* central nervous system (CNS). We found that Nerfin-1 inactivation causes tumorigenic phenotypes in the central brain (CB) and the ventral nerve cord (VNC) but not the optic lobes (OLs). In contrast, Lola inactivation leads to tumour overgrowth specifically in the OLs. We identify Chinmo, a temporal transcription factor, and its regulation by ecdysone signalling as key determinants of the oncogenic competence in different regions of the brain, influencing the tumorigenic outcome of dedifferentiation. This work provides a fundamental framework to understand how oncogenic competence arises beyond genetic mutations.

**Significance statement:** In the CNS, the same tumorigenic mutation has been shown cause differential oncogenic outcomes in different regions of the brain. The mechanism underlying this phenomenon remains largely unknown. We have previously demonstrated that malignant brain tumours can be induced via neuronal dedifferentiation in the *Drosophila* CNS. Here, we demonstrate that dedifferentiated neural stem cells drive tumorigenesis in a region-specific manner, accounted for by region-specific expression and regulation of temporal factors by cell-intrinsic and hormonal signals. Together, this work extends our understandings of how brain regionalisation can be a constraint to oncogenesis.

## Introduction

Genetic mutations are not the sole determinants of malignant cancer outcomes. In fact, individuals can accumulate mutations with oncogenic potential, yet most are eliminated without leading to malignancy. Oncogenic competence is defined by a combination of genetic mutations, lineage-specific factors, and microenvironmental cues (1). Many brain and central nervous system (CNS) cancers originate from excessively proliferative neural stem-progenitor cells during embryonic development (2, 3). Perturbations in signalling pathways and transcription factors involved in neural development can initiate tumour overgrowth in the CNS (4). Interestingly, distinct oncogenic mutations are associated with specific anatomical regions of the CNS (5–7), suggesting that oncogenic competence is regionally regulated. However, the mechanisms by which region-specific cues influence cancerous transformation remain poorly understood.

The *Drosophila* CNS is a well-established model for studying neurogenesis, cell fate specification, and maintenance during development. In *Drosophila*, neural stem cell-like progenitors, known as neuroblasts (NBs), generate neurons and glia across all regions of the CNS. Here, the CNS is broadly divided into three main regions: the ventral nerve cord (VNC), the central brain (CB), and the optic lobes (OLs) (Fig. 1A). From embryonic to mid-pupal development, NBs undergo a limited number of divisions until their termination, by which the mechanism differs between regions (8–10). In the CB/VNC, NB termination mainly occurs via terminal symmetric division mediated by the homeobox transcription factor Prospero (Pros) (11). Whereas in the optic lobe (OL), medulla NBs undergo termination through a combination of apoptosis, terminal symmetric division, and gliogenic switch (12). Thus, NBs terminate in a region-specific fashion to meet the differential demands of neuronal function during adulthood.

**Figure 1:**
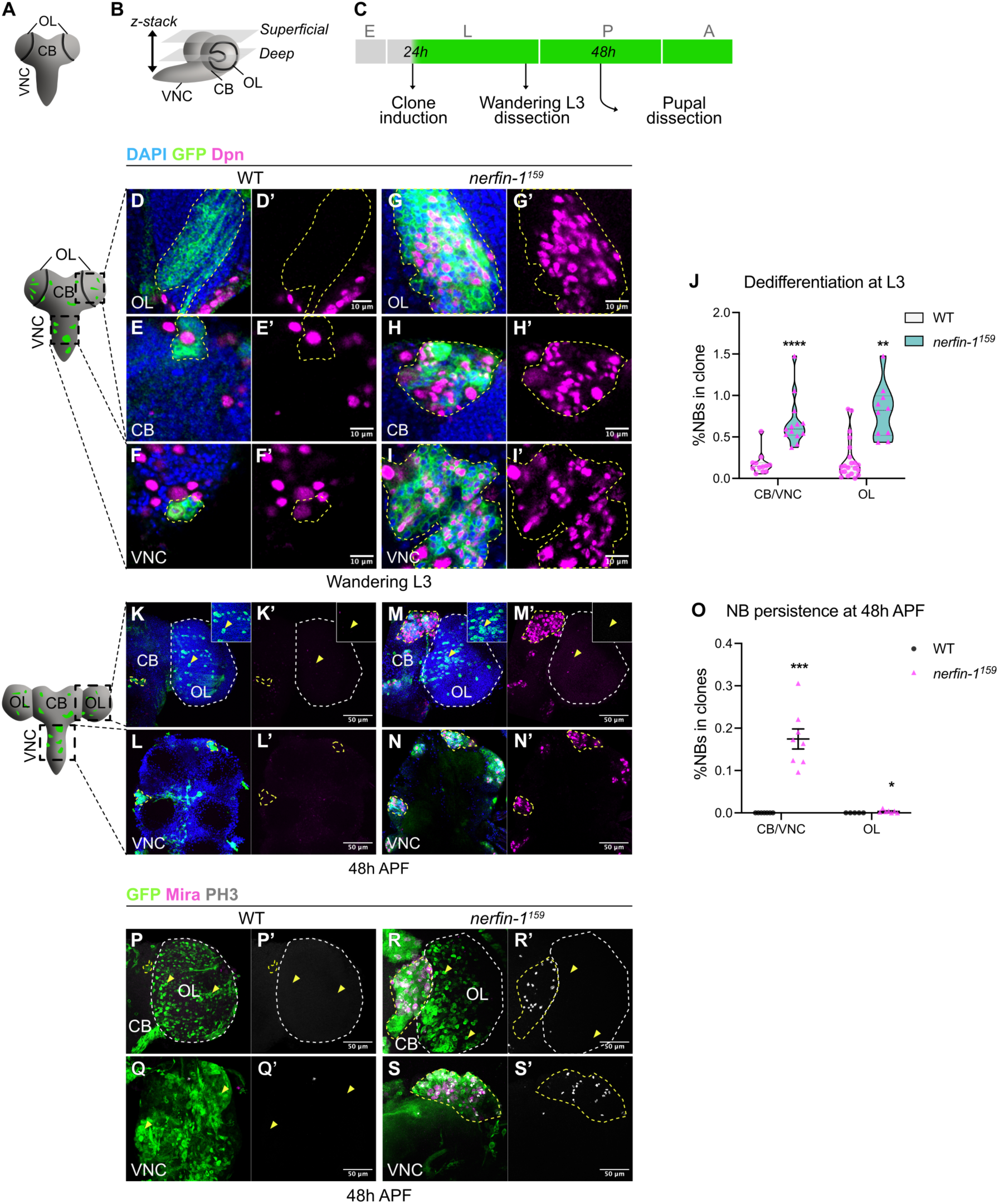
*nerfin-1^159^* dedifferentiated NBs are selectively eliminated in the pupal OL. **A-B.** Schematic depicting **(A)** the frontal view and **(B)** the side view of the larval CNS that consists of the central brain (CB), the ventral nerve cord (VNC) and the optic lobes (OLs). Representative confocal images of the larval brain are taken from superficial or deep layers. **C.** Schematic summarising our experimental procedure. **D-I’**. Single confocal sections in the deep layers of **(D-F’)** WT or **(G-I’)** *nerfin-1^159^* clones (dashed lines) in **(D-D’, G-G’)** the OL, **(E-E’, H-H’)** CB, and **(F-F’, I-I’)** VNC at wandering L3 stage. NBs marked by Dpn (magenta), GFP (green), DAPI (blue). Scale bars: 10 μm. **J.** Quantification of the percentages of NBs in WT and *nerfin-1^159^* clones in the CB/VNC and OL at wandering L3 stage. n=12, 10, 9, and 10 clones. **K-N’.** Maximum projections of **(K-K’, M-M’)** the brain lobes (CB and OL, separated by white dash lines) and **(L-L’, N-N’)** VNC with **(K-L’)** WT or **(M-N’)** *nerfin-1^159^* clones (yellow dash lines) at 48h APF. DAPI (blue), GFP (green), Dpn (magenta). Arrowheads indicate GFP^+^ cells. Scale bars: 10 μm. **O.** Quantification of the percentages of NBs in WT and *nerfin-1^159^*clones in the CB/VNC and OL at 48h APF. n=7, 8, 5, and 7. **P-S’.** Maximum projections of **(P-P’, R-R’)** the brain lobes (CB and OL, separated by white dash lines) and **(Q-Q’, S-S’)** VNC with **(P-Q’)** WT or **(R-S’)** *nerfin-1^159^* clones at 48h APF. Arrowheads indicate GFP^+^ cells. Yellow dash lines indicate tumour formation in the CB/VNC. Mature NBs are marked by Miranda (Mira, magenta), mitotic cells are marked by Phospho-Histone 3 (PH3, grey), GFP (green). Scale bars: 50 μm. Data information: Data are represented as mean ± SEM. *p*-values were obtained using Mann-Whitney test. \*\*\*\**p* < 0.0001, \*\*\**p* < 0.001, \*\**p* < 0.005, \**p* < 0.05.

Recent studies have shown that temporal transcription factors (tTFs) and their target genes regulate NB termination in the CB and the VNC. These tTFs are sequentially expressed in NBs as they age, not only specifying distinct neuronal identities in their progeny but also influencing cell cycle speed and the timing of NB termination. For instance, during early larval development, NBs in the CB/VNC express early temporal factors: IGF-II mRNA-binding proteins (Imp) and the BTB-zinc finger transcription factors Chronologically inappropriate morphogenesis (Chinmo) (11, 13, 14). As the animal matures, NBs shift to express late temporal factors, namely Syncrip (Syp) and Broad (Br) (11, 13, 14). Imp and Syp modulate NB proliferation through opposing gradients, with Imp promoting the proliferation whereas Syp driving the termination of NBs (14, 15). Thus, an extended period of Imp expression or inhibition of Syp can prolong NB proliferation (15). Temporal patterning is also observed in the OL medulla progenitors, the NE in which Imp and Chinmo define the early temporal window, while Syp, Br, and Eip93F define the late temporal window (16–18). However, it is unclear if this temporal cascade influences medulla NB termination.

In this study, we investigate how dedifferentiation - triggered by the loss of neuronal cell fate maintenance – can lead to contrasting tumorigenic outcomes. Previously, we and others have shown that neuronal cell fate requires active maintenance by various transcription factors, such as the zinc finger transcription factor Nervous finger 1 (Nerfin-1) and the BTB-zinc finger transcription factor Longitudinals lacking (Lola) (19–22). Failure to maintain the neuronal identity can result in the formation of supernumerary NB that can drive tumour overgrowth. Here, we demonstrate that oncogenic competence of dedifferentiated NBs is region-specific. In the CB and VNC, the loss of Nerfin-1 function leads to malignant tumour overgrowth, with ectopic NBs continuing to proliferate beyond the normal neurogenesis period (21). In contrast, in the OLs, we show that ectopic NBs are eliminated through cell death and differentiation. Unlike Nerfin-1, Lola loss-of-function elicits tumorigenesis in the OL (20). Mechanistically, we identified the early tTF Chinmo as a key regulator of this differential response and that the cell’s ability to respond to Chinmo is mediated through ecdysone signalling. As such, increasing Chinmo levels confers oncogenic competence to dedifferentiated NBs induced by Nerfin-1 inactivation in the OL. Conversely, reducing Chinmo levels in those in the CB/VNC or in *lola^-/-^* clones promote elimination of dedifferentiated NBs. These findings highlight that the differential expression and regulation of oncogenic tTFs such as Chinmo, play a crucial role in determining tumorigenic outcome in response to dedifferentiation triggers. Overall, our study shed light on mechanisms of oncogenic competency and offers insights into how certain regions of the brain are more susceptible to tumorigenesis than others.

## Results

### *nerfin-1* loss-of-function (*nerfin-1^159^)* causes malignant tumour overgrowth in the CB/VNC but not the OL

We and others have previously reported that *nerfin-1* loss-of-function leads to the dedifferentiation of neurons into ectopic NBs (19, 21, 22). In this study, to investigate region-specific regulation of dedifferentiation, we began by inducing *nerfin-1^159^*-null mutant (23) loss-of-function MARCM clones at 24 hrs ALH (After Larval Hatching) in all the regions of the larval CNS (CB, VNC, and OLs) (Fig. 1A-C). At wandering L3 stage, control wildtype (WT) MARCM clones in the OL often consist of a few NBs, marked by pan-NB marker Deadpan (Dpn) on the superficial layers and none in the deep layers where neurons reside (Fig. 1D-D’). Whereas *nerfin-1^159^* clones had many ectopic Dpn^+^ NBs in the deep layers (Fig. 1G-G’, J). Similarly, on the superficial layers of the ventral side of the CB (where type I but not type II NBs reside) and the VNC, WT clones consisted of one NB and none in the deep layers that are occupied by neurons (Fig. 1E-F’). In contrast, many ectopic NBs were observed in the deep layers of *nerfin-1^159^* clones in the CB/VNC (Fig. 1H-J). These data indicate that neurons revert into ectopic NBs in all regions of the CNS upon Nerfin-1 inactivation.

In the developing CB/VNC, NBs terminally differentiate at around 24 hrs after pupal formation (APF) (11), and the medulla NBs of the OL terminate between 12-24 hrs APF (12). A key characteristic of malignant tumours is their failure to obey signals to terminally differentiate and their ability to proliferate beyond the normal period of neurogenesis. To examine these characteristics, we analysed the brains at 48 hrs APF. In all regions of the brain, no NB persisted in WT clones (Fig. 1K-L’, O). However, we observed a discrepancy between the CB/VNC and the OL that contain *nerfin-1^159^* clones. In the CB/VNC, we found that *nerfin-1^159^* clones were enriched with ectopic NBs (Fig. 1M-O). Whereas in the OLs, *nerfin-1^159^* clones became dispersed and devoid of Dpn expression (Fig. 1M-M’, O), indicative of neuronal differentiation and migration (24).

To ascertain whether *nerfin-1^159^* NBs undergo proliferation beyond the normal period of neurogenesis, we stained the pupal brains at 48 hrs APF for proliferative NBs marked by Miranda (Mira) and Phospho-Histone 3 (PH3). In all regions of the CNS, WT clones did not exhibit any Mira^+^ PH3^+^ persistent NBs at 48 hrs APF (Fig. 1P-Q’). In contrast, we found that Mira^+^ PH3^+^ NBs were present in *nerfin-1^159^* clones in the CB/ VNC (Fig. 1R-S’, yellow dash lines) but absent in clones in the OL (Fig. 1R-R’, arrowheads). Collectively, our results reveal that the same genetic mutation (*nerfin-1^159^*) can cause differential consequences in a region-specific manner: dedifferentiated NBs drive tumorigenic overgrowth in the CB/VNC but are eliminated in the OL.

### *nerfin-1^159^* dedifferentiated NBs in the OL are eliminated via a combination of Pros-mediated differentiation and apoptosis-mediated cell death

To test how *nerfin-1^159^* dedifferentiated NBs become eliminated in the pupal OL, we examined whether these NBs undergo differentiation or a gliogenic switch – mechanisms responsible for WT OL NB termination (12). First, we assayed for the expression of the neuronal marker Elav and glial marker Repo in *nerfin-1^159^* clones at 48 hrs APF. We found that *nerfin-1^159^* clones contained Elav^+^ neurons (Fig. 2A-B’’, cyan arrowheads) and Repo^+^ glial cells (Fig. 2A-B’’, magenta arrowheads) in both pupal OL and VNC. Additionally, the rate of neuronal differentiation (percentage of Elav^+^ volume per region of interest, see Material and Methods) was significantly higher in the OL than the VNC at 48 hrs APF (Fig. 2C). The terminal differentiation of WT NBs via symmetric division is mediated by the homeobox transcription factor Pros (11, 12). At 24 hrs APF, we found that *nerfin-1^159^* NBs in both the OL and VNC expressed low levels of Pros (Fig. 2D-E’’, arrowheads), suggesting that the elimination of these ectopic NBs requires terminal differentiation. To address this, we knocked down Pros in *nerfin-1^159^* clones. At wandering L3 stage, ectopic NBs were found in both *pros^RNAi^* and *pros^RNAi^;nerfin-1^159^* clones in the deep layers of the OLs (Fig. 2F-H). By 48 hrs APF, *pros^RNAi^;nerfin-1^159^*clones exhibited significantly increased NB persistence compared to *nerfin-1^159^*or *pros^RNAi^* clones (Fig. 2I-K’, M). This suggests that Pros-mediated terminal differentiation contributes to the elimination of *nerfin-1^159^*dedifferentiated NBs in the pupal OL. To elucidate whether the gliogenic switch also contributes to the elimination of *nerfin-1^159^* dedifferentiated NBs, we knocked down the glial cell fate determinant Glial-cell-missing (Gcm) by RNAi. However, this manipulation did not significantly increase the persistence of ectopic NBs at 48 hrs APF (Fig. 2L-L’, N). Hence, an acquisition of the glial cell fate does not substantial contribute to the elimination of *nerfin-1^159^* dedifferentiated NBs in the pupal OL. A subset of NBs in the developing OL terminates via apoptosis-mediated cell death during early pupal development (12). At wandering L3 stage, we found that some dedifferentiated NBs in *nerfin-1^159^* clones expressed the apoptotic marker Caspase-1 (Dcp-1) (Fig. S1A-A’, arrowheads). To address if apoptosis is required for the elimination of *nerfin-1^159^* dedifferentiated NBs, we inhibited apoptosis by overexpressing the apoptotic inhibitor p35 in *nerfin-1^159^* clones. This perturbation resulted in the elimination of Dcp-1 expression in the NBs at wandering L3 stage, indicating that apoptosis was suppressed (Fig. S1B-B’). This was accompanied by a small increase in the NB persistence in *nerfin-1^159^* clones at 48 hrs APF (fig. S1C-E). However, most of the *UAS-p35;nerfin-1^159^*cells in the OLs were devoid of Dpn (Fig. S1D-D’, white arrowheads). We found that many *UAS-p35;nerfin-1^159^* cells in the pupal OL expressed either Elav or Repo (Fig. S1F-F’’), suggesting that the cells that do not die may still undergo terminal differentiation or acquire a glial cell fate. Together, our data suggest that the elimination of *nerfin-1^159^*dedifferentiated NBs in the pupal OL involves a combination of Pros-mediated terminal differentiation and apoptosis.

**Figure 2:**
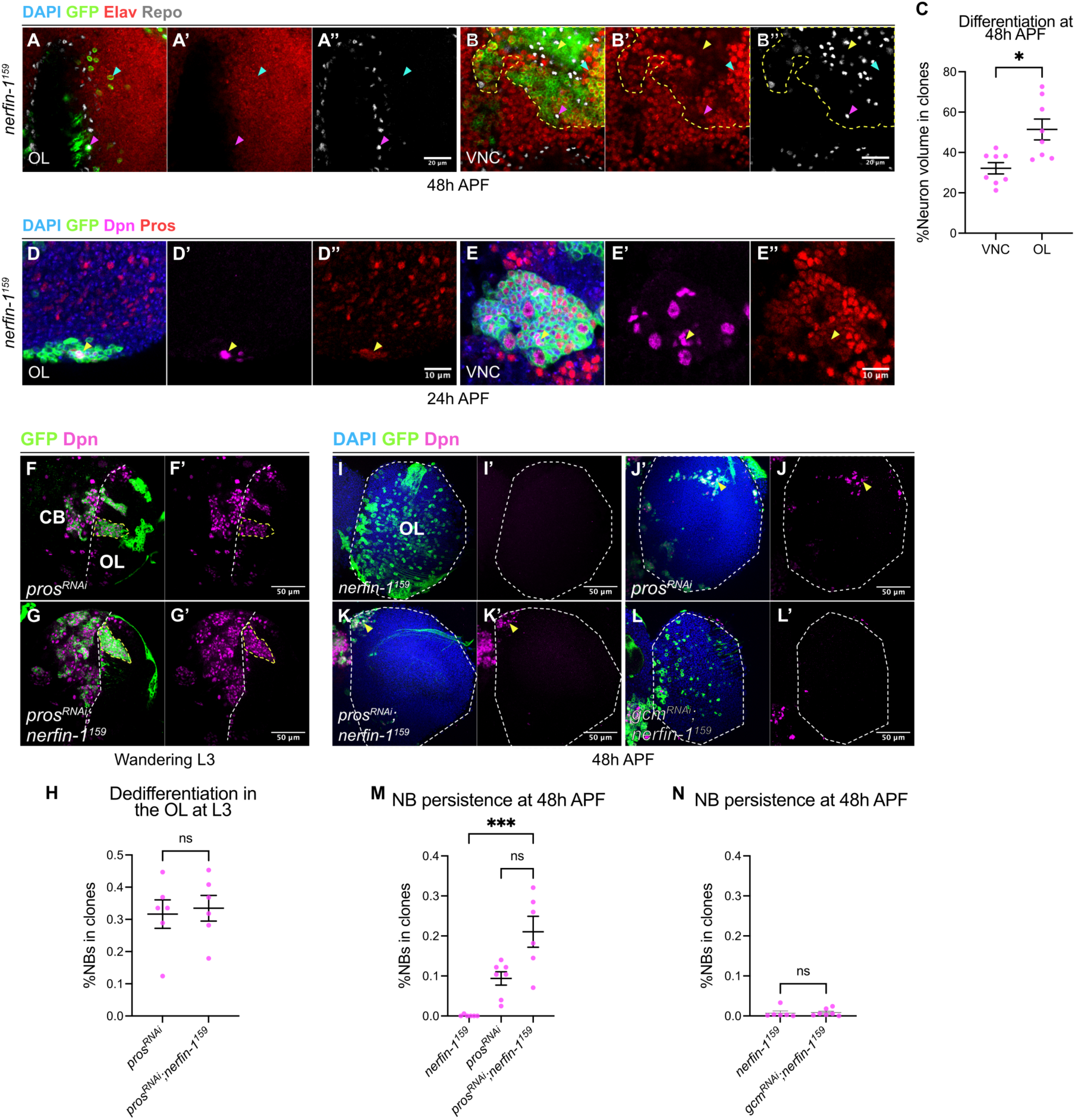
*nerfin-1^159^*dedifferentiated NBs in the OL are eliminated via Pros-mediated terminal differentiation during pupal development. **A-B’’.** Single confocal sections of *nerfin-1^159^* clones in **(A-A’’)** the OL and **(B-B’’)** the VNC at 48h APF. GFP (green), Elav (red), Repo (grey). In the VNC, yellow dash lines outline the mutant clone. Yellow arrowhead indicates Elav^-^Repo^-^ mutant cell. Cyan and magenta arrowheads indicate Elav^+^Repo^-^ and Elav^-^Repo^+^ mutant cells, respectively. Scale bars: 20 μm. **C.** Quantification of the percentages of the volume of neurons in *nerfin-1^159^*clones in the OL and the VNC at 48h APF. VNC: n=8, and 8. **D-E’’.** Single confocal sections of *nerfin-1^159^* clones in **(D-D’’)** the OL and **(E-E’’)** the VNC at 24h APF. DAPI (blue), GFP (green), Dpn (magenta), Pros (red). Scale bars: 10 μm. **F-G’.** Single confocal sections in the deep layers of the brain lobes at wandering L3 stage with **(F-F’)** *pros^RNAi^* and **(G-G’)** *pros^RNAi^;nerfin-1^159^* clones (yellow dash lines). GFP (green), Dpn (magenta). White dash lines outline the CB/OL borders. Scale bars: 50 μm. **H.** Quantification NB percentages in *pros^RNAi^* and *pros^RNAi^;nerfin-1^159^* clones in the OL at wandering L3 stage. n=6, and 6 clones. **I-L’.** Maximum projections of the OL (dash lines) at 48h APF with **(I-I’)** *nerfin-1^159^*, **(J-J’)** *pros^RNAi^*, **(K-K’)** *pros^RNAi^;nerfin-1^159^* and **(L-L’)** *gcm^RNAi^;nerfin-1^159^*clones. Arrowheads indicate ectopic NBs. DAPI (blue), GFP (green), Dpn (magenta). Scale bars: 50 μm. **M.** Quantifications of NB percentages in *nerfin-1^159^*, *pros^RNAi^*, and *pros^RNAi^;nerfin-1^159^* clones in the OLs at 48h APF. n=6, 6, and 7. **N.** Quantifications of the percentages of NBs in *nerfin-1^159^* and *gcm^RNAi^;nerfin-1^159^* clones in the OLs at 48h APF. n=6, and 7. Data information: Data are represented as mean ± SEM. *p*-values were obtained using Mann-Whitney test and Kruskal-Wallis test with Dunn’s test to correct for multiple comparisons. \*\*\*\**p* < 0.0001, \**p* < 0.05.

### The elimination *of nerfin-1^159^* dedifferentiated NBs in the pupal OL occurs independently of Myc

In the VNC, we previously showed that *nerfin-1^159^-*induced neuronal dedifferentiation is dependent on the expression of the growth regulator Myc (21). Interestingly, our immunostaining showed that Myc was expressed in *nerfin-1^159^* clones in the CB and VNC but not in the OL at wandering L3 stage (Fig. S1G-H’). Hence, it is plausible the elimination of *nerfin-1^159^* dedifferentiated NBs in the pupal OL is caused by the lack of Myc. To test this hypothesis, we overexpressed Myc in *nerfin-1^159^* clones. While this manipulation was sufficient to increase Myc levels in *nerfin-1^159^* clones in the OL, it was not sufficient to promote the persistence of ectopic NBs at 48 hrs APF (Fig. S1E, I-J’’).

### The elimination *of nerfin-1^159^* dedifferentiated NBs in the pupal OL is due to the absence of the oncogenic temporal factor Chinmo

In the VNC, the malignant tumour competency is determined by the expression of the oncogenic tTF Chinmo, such that cells generated during the Imp^+^Chinmo^+^ window are susceptible to malignant tumour overgrowth (25, 26). Whereas late-born cells from the Syp^+^Br^+^ temporal window have the propensity to undergo terminal differentiation. To investigate if Chinmo is involved in the elimination of *nerfin-1^159^* dedifferentiated NBs in the pupal OL, we first examined Chinmo expression in *nerfin-1^159^* clones at wandering L3 stage. Consistent with previous reports (26), we found that Chinmo was expressed in many WT neurons as well as *nerfin-1^159^* dedifferentiated NBs in the CB and VNC (Fig. 3A-B’’ and Fig. S2A-A’’). In contrast, the majority of neurons and *nerfin-1^159^* dedifferentiated NBs in the OL did not express Chinmo (Fig. 3C-C’’ and Fig. S2B-B’’).

**Figure 3:**
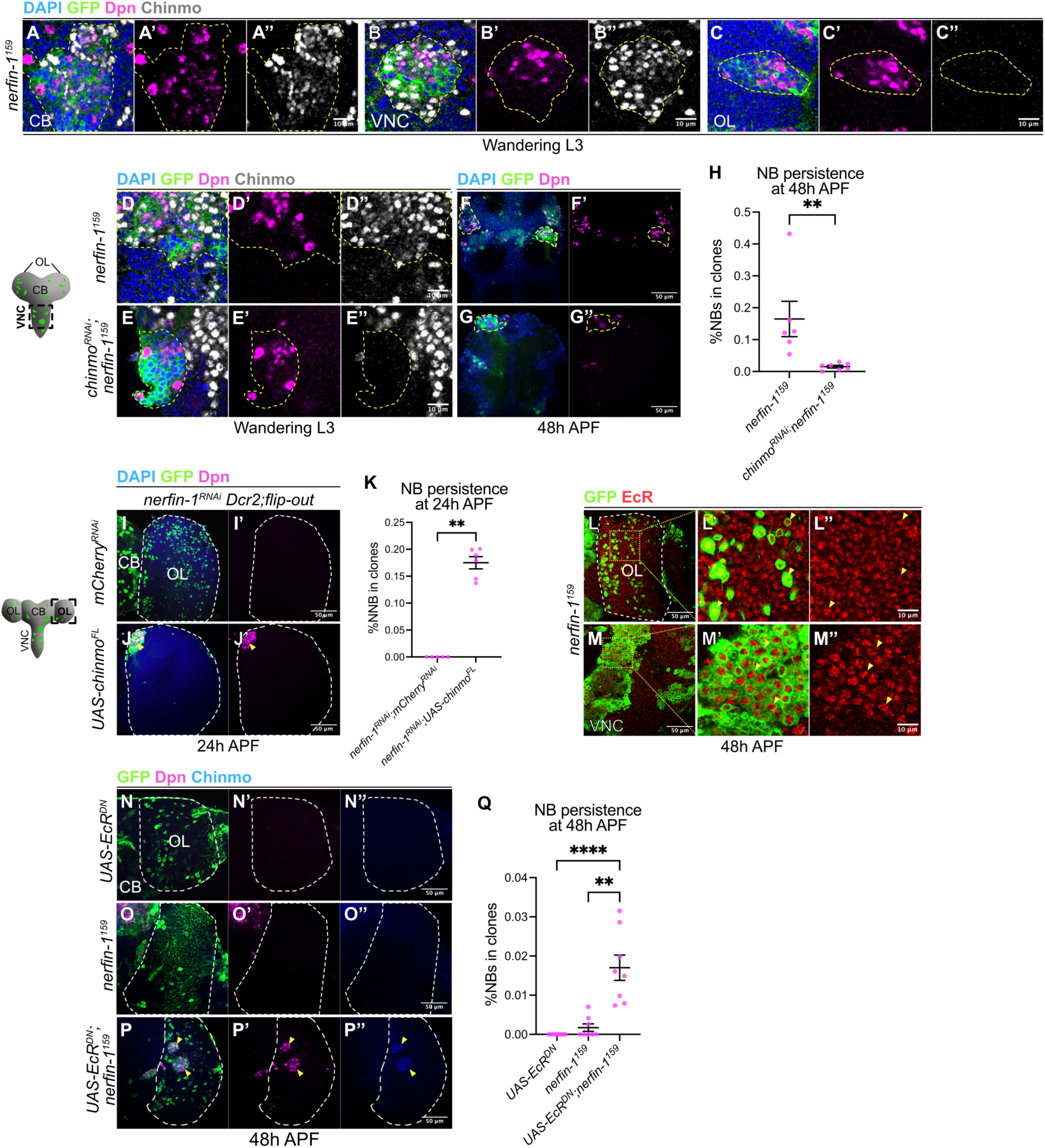
Chinmo promotes the persistence of *nerfin-1^159^* ectopic NBs in the pupal CNS. **A-C’’.** Single confocal sections of *nerfin-1^159^* clones (dash lines) in **(A-A’’)** the CB, **(B-B’’)** VNC and **(C-C’’)** OL at wandering L3 stage. DAPI (blue), GFP (green), Dpn (magenta), Chinmo (grey). Scale bars: 10 μm. **D-E’’** Single confocal sections of **(D-D’’)** *nerfin-1^159^* and **(E-E’’)** *chinmo^RNAi^*;*nerfin-1^159^* clones (dash lines) in the VNC at wandering L3 stage. DAPI (blue), GFP (green), Dpn (magenta), Chinmo (grey). Scale bars: 10 μm. **F-G’.** Maximum projections of the VNC with **(F-F’)** *nerfin-1^159^* and **(G-G’)** *chinmo^RNAi^*;*nerfin-1^159^* clones (dash lines) at 48h APF. DAPI (blue), GFP (green), Dpn (magenta). Scale bars: 50 μm. **H.** Quantification of the percentages of NBs in *nerfin-1^159^* and *chinmo^RNAi^;nerfin-1^159^*clones in the VNC at 48h APF. n=6, and 7. **I-J’.** Maximum projections of the OLs (dash lines) at 24h APF with **(I-I’)** *nerfin-1^RNAi^;mCherry^RNAi^* and **(J-J’)** *nerfin-1^RNAi^;UAS-chinmo^FL^*flip-out clones. Arrowheads indicate persistent ectopic NBs. DAPI (blue), GFP (green), Dpn (magenta). Scale bars: 50 μm. **K.** Quantification of the percentages of NBs in *nerfin-1^RNAi^;mCherry^RNAi^* and *nerfin-1^RNAi^;UAS-chinmo^FL^* flip-out clones in the OLs at 24h APF. n=5, and 6. **L-M’’.** Left panels: single confocal section of the **(L-L’’)** OL (dash lines) and **(M-M’’)** VNC at 48h APF with *nerfin-1^159^* clones (arrowheads). Right panels are boxed area in left panels. GFP (green), EcR (red). Scale bars: 50 and 10 μm. **N-P’’** Single confocal sections of the OLs (dash lines) at 48h APF with **(N-N’’)** *UAS-EcR^DN^*, **(O-O’’)** *nerfin-1^159^* and **(P-P’’)** *UAS-EcR^DN^*;*nerfin-1^159^*clones. Arrowheads indicate persistent ectopic Chinmo^+^ NBs. GFP (green), Dpn (magenta), Chinmo (blue). Scale bars: 50 μm. **Q.** Quantification of the percentages of NBs in *UAS-EcR^DN^, nerfin-1^159^*, and *UAS-EcR^DN^;nerfin-1^159^* clones in the OLs at 48h APF. n=8, 8, and 8. Data information: Data are represented as mean ± SEM. *p*-values were obtained using Mann-Whitney test and Kruskal-Wallis test with Dunn’s test to correct for multiple comparisons. \*\*\*\**p* < 0.0001, \*\**p* < 0.005.

To test whether Chinmo is functionally required to promote the persistence of dedifferentiated NBs in the CB/VNC, we expressed *chinmo^RNAi^* (see Materials and methods for details of RNAi construction) in *nerfin-1^159^* clones. At wandering L3 stage, the expression of *chinmo^RNAi^* effectively reduced Chinmo levels in *nerfin-1^159^* clones in the VNC (Fig. 3D-E’’). At 48 hrs APF, *chinmo^RNAi^;nerfin-1^159^* clones significantly reduced NB persistence compared to *nerfin-1^159^* clones (Fig. 3F-H). Hence, Chinmo is required to promote tumorigenesis of *nerfin-1^159^*dedifferentiated NBs in the pupal CB/VNC.

Next, we sought to address whether the absence of Chinmo accounts for the elimination of *nerfin-1^159^*dedifferentiated NBs in the pupal OL. To do so, we overexpressed a full-length version of *chinmo* (*chinmo^FL^*) (27) in *nerfin-1^RNAi^* flip-out clones. By 24 hrs APF, *nerfin-1^RNAi^;mCherry^RNAi^*control clones in the OL were devoid of Dpn and cells were dispersed across the OLs (Fig. 3I-I’). In contrast, *nerfin-1^RNAi^*;*chinmo^FL^* clones contained persistent ectopic NBs (Fig. 3J-K). From mid-larval stages, medulla NBs located on the cortical surface are generated from a pseudostratified neuroepithelium (NE) of the OL named the NE-NB transition (28, 29). Importantly, Chinmo was previously shown to play a critical role in regulating the NE-NB transition (17). In early larval development, Chinmo is expressed in the NE to promote NE self-renewal whereas from mid-larval stages, Chinmo is downregulated, allowing NE differentiation into NBs. Thus, the overexpression of Chinmo in the NE can result in a delayed NE-NB transition which can subsequently delays NB termination (17). To overexpress Chinmo in a subset of medulla NB (and not NE), we utilised *tub-GAL80^ts^*;*eyR16F10-GAL4* (12) to drive the expression of *chinmo^FL^* temporally from early L3. At wandering L3 stage, ectopic NBs were found in the deep layers of the medulla upon the induction of *nerfin-1^RNAi^* (Fig. S2C-E’). At 24 hrs APF, *nerfin-1^RNAi^;mCherry^RNAi^*and *lacZ^RNAi^;mCherry^RNAi^* OLs exhibited no persistent NBs (fig. S2F-G’, I). In contrast, many ectopic NBs were observed in the OLs upon the misexpression of *nerfin-1^RNAi^;UAS-chinmo^FL^* (Fig. S2H-I). Altogether, our data suggest that Chinmo is sufficient to prevent the elimination of *nerfin-1* loss-of-function NBs in the pupal OL.

### Chinmo is repressed by ecdysone signalling in *nerfin-1^159^* clones in the OL

We next investigated how Chinmo expression is regulated in *nerfin-1^159^* clones in the OL. In the developing OL medulla, Chinmo has been shown to be expressed from early to mid-larval stages (16, 17). As *nerfin-1^159^* clones in the larval OL were devoid of Chinmo expression (Fig. 3C-C’’), we next investigated how Chinmo expression is repressed, and whether this is regulated by transcriptional or post-transcriptional mechanisms. We first assessed the expression of *chinmo*-lacZ transcriptional reporter in *nerfin-1^159^*clones in the CB, VNC and the OL at wandering L3 stage. In the CB and VNC, *chinmo*-lacZ was expressed in *nerfin-1^159^* clones (Fig. S2J-K’’). On the contrary, *chinmo*-lacZ was absent in *nerfin-1^159^* clones in the OL (Fig. S2J’’), indicating that *chinmo* is transcriptionally silenced in *nerfin-1^159^* clones the OL but not the CB/VNC. We then also tested if *chinmo* can be post-transcriptionally controlled in *nerfin-1^159^* clones. To do so, we utilised a reporter in which the mCherry coding sequence is flanked by the 5’ and 3’ UTRs of *chinmo* (*UAS-mCherry-chinmo^UTRs^*) (17). If *chinmo* is regulated post-transcriptionally, it is expected that clones would not express mCherry. However, we found that similar to WT clones *nerfin-1^RNAi^*flip-out clones expressed mCherry (Fig. S2L-M’’, dash lines), indicating that Chinmo expression is regulated via transcriptional rather than post-transcriptional mechanisms.

Ecdysone is a steroid hormone and a central regulator of developmental transitions in insects including *Drosophila* (30). In the WT CNS, it has been previously shown that ecdysone signalling at the critical weight (mid-L3 stage) is required to repress Chinmo transcription specifically in the OL but not the CB/VNC (17). Therefore, it is plausible that ecdysone signalling inhibits Chinmo expression in *nerfin-1^159^*dedifferentiated NBs in the OL. We found that Ecdysone Receptor (EcR) was expressed in *nerfin-1^159^* clones at a similar level to the surrounding WT cells at 48 hrs APF in the OL and VNC (Fig. 3L-M’’, arrowheads), suggesting that *nerfin-1^159^* clones may be responsive to ecdysone signals. To inhibit ecdysone signalling, we expressed a dominant negative form of EcR (EcR^DN^) (31) in *nerfin-1^159^* clones and assayed for Chinmo expression at 48 hrs APF. While *UAS-EcR^DN^* expression alone was not sufficient to upregulate Chinmo, nor promote the persistence of NBs (Fig. 3N-N’’,Q), *UAS-EcR^DN^* expression in *nerfin-1^159^* clones caused Chinmo to be upregulated and resulted in increased NB persistence in the OL (Fig. 3O-Q). These data suggest that ecdysone signalling is necessary for Chinmo silencing in *nerfin-1^159^*clones in the pupal OL, causing the dedifferentiated NBs to lose tumorigenicity and differentiate. It is of note that *nerfin-1^159^* clones in the VNC also expressed EcR (Fig. 3M-M’’). Nevertheless, because *nerfin-1^159^* dedifferentiated NBs persist in the pupal CB/VNC, it is likely that they do not respond to ecdysone signals.

We also examined if known post-transcriptional regulators of Chinmo such as the RNA-binding protein Imp (32) or the microRNA *let-7* (33) can upregulate Chinmo expression in *nerfin-1^159^*clones and thereby, promote ectopic NB persistence in the pupal OL. At 24 hrs APF, we found that the overexpression of Imp in *nerfin-1^159^* clones was able to downregulate its antagonist Syp but was however not sufficient to upregulate Chinmo expression (Fig. S2N-O’’’). Additionally, by 48 hrs APF, the percentages of persistent NBs in *UAS-Imp;;nerfin-1^159^* clones were comparable to that of *nerfin-1^159^* clones (Fig. S2P). As such, Imp is not sufficient to upregulate Chinmo expression or promote tumour overgrowth in the OL. To functionally test the role of *let-7*, we misexpressed *let-7^sponge^* in *nerfin-1^159^* clones and assayed for ectopic NBs in the OL at 48 hrs APF. Similar to Imp overexpression, no NB persisted in both *let-7^sponge^;nerfin-1^159^*(n=6/6 lobes) and *nerfin-1^159^* clones (n=3/3 lobes) (Fig. S2Q-R’), suggesting that *let-7* is unable to promote dedifferentiated NBs to persist in the pupal OL. Altogether, our results are consistent with the notion that *chinmo* is transcriptionally silenced by ecdysone signalling between larval-pupal development in *nerfin-1^159^* clones in the OL. Hence, altering post-transcriptional modulators of Chinmo is not sufficient to prevent the elimination of *nerfin-1^159^* dedifferentiated NBs.

### Lola and Nerfin-1 have different expression patterns and regulate different gene sets in the developing OL medulla

*lola* loss-of-function (*lola^E76^* – a null mutation where all Lola isoforms are deleted) (Goeke et al., 2003) causes neuronal dedifferentiation and consequently, tumour formation specifically in the adult OL (Southall et al., 2014). Indeed, we observed ectopic NB formation in *lola^E76^* clones in the deep layers of the OL at wandering L3 stage (Fig. 4A-B), indicating neuronal dedifferentiation. At 48 hrs APF, unlike WT clones in which NBs already terminated, *lola^E76^*clones in the OL consisted of multiple ectopic Dpn^+^ NBs (Fig. 4C-D) as well as many Mira^+^PH3^+^ NBs (Fig. 4E-F). Therefore, it is likely that these NBs were capable of proliferation beyond the normal period of neurogenesis. Notably, we found that some cells within *lola^E76^* clones were able to undergo differentiation, as indicated by the presence of neuronal marker Elav and glial marker Repo (Fig. S3A-A’’’). Furthermore, nuclear expression of Pros was detected in *lola^E76^* NBs similar to WT cells in the OL at 24 hrs APF (Fig. S1B-C’’), suggesting that *lola^E76^* ectopic NBs can undergo Pros-mediated terminal symmetric division. Together, these data suggest that beyond region-specific mechanisms, the ability to form tumours is also determined by the nature of the genetic mutations; while *nerfin-1^159^* dedifferentiated NBs are eliminated, *lola^E76^* dedifferentiated NBs remain proliferative to initiate tumorigenesis in the pupal OL.

**Figure 4:**
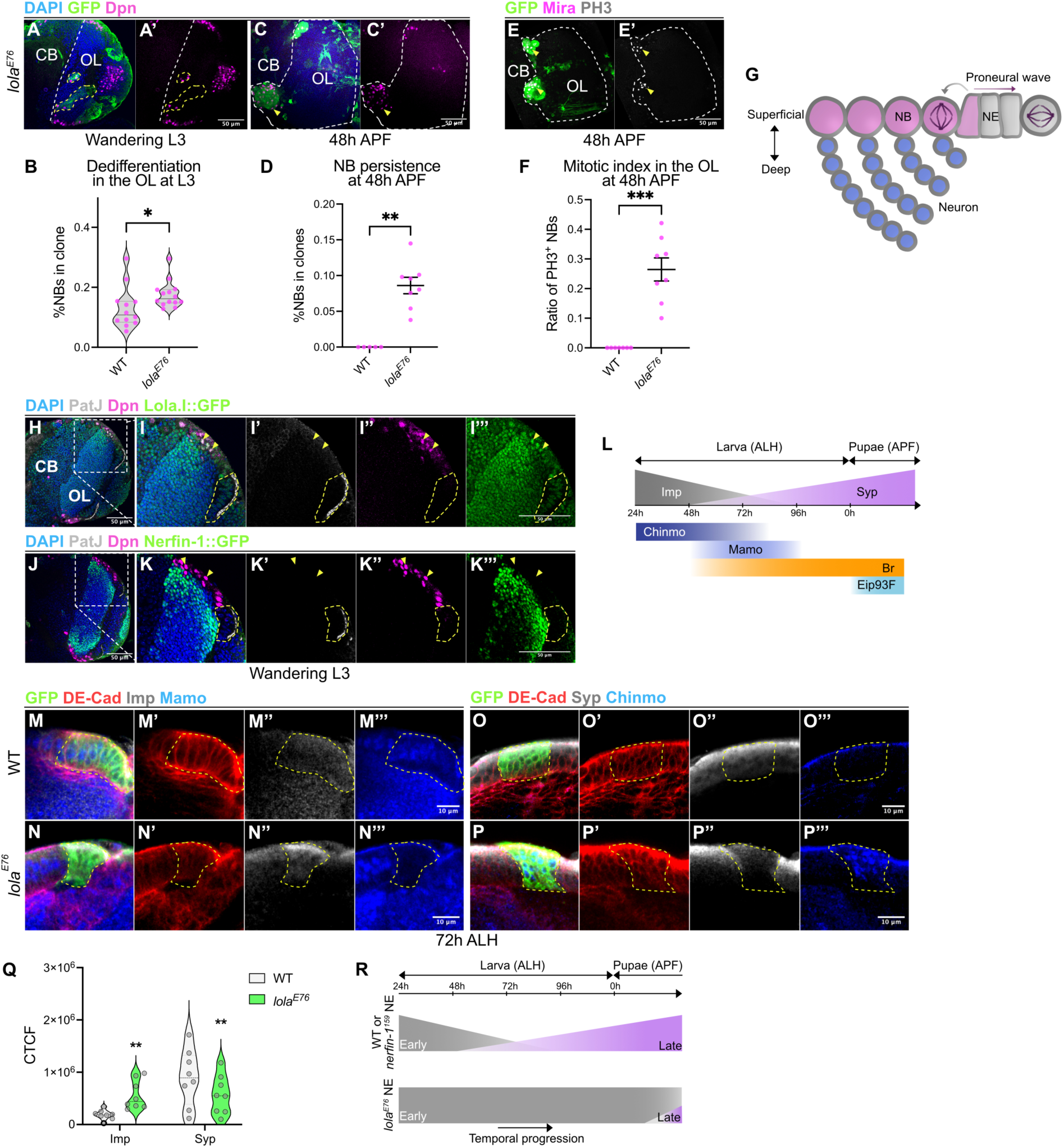
Lola promotes temporal progression of the larval NE. **A-A’.** Single section of the late L3 OL (deep layer) with *lola^E76^* clones (dash lines). DAPI (blue), GFP (green), Dpn (magenta). Scale bars: 50 μm. **B-B’.** Quantification of NB percentages in WT and *lola^E76^* clones in the late L3 OLs. n=12 and 14 clones. **C-C’.** Maximum projections of the OL (dash lines) at 48h APF with *lola^E76^* clones (arrowheads). DAPI (blue), GFP (green), Dpn (magenta). Scale bars: 50 μm. **D.** Quantification of NB percentages in WT and *lola^E76^* clones in the OLs at 48h APF. n=5 and 8. **E-E’.** Maximum projections of the OL (dash lines) at 48h APF with *lola^E76^* clones (arrowheads). GFP (green), Mira (magenta), PH3 (grey). Scale bars: 50 μm. **F.** Quantification of mitotic NB percentages in WT and *lola^E76^* clones in the OLs at 48h APF. n=7 and 8. **G.** Schematic depicting the larval OL medulla. **H-K’’’** Single sections of late L3 brain lobes expressing **(H-I’’’)** Lola.I::GFP and **(J-K’’’)** Nerfin-1::GFP. **(I-I’’’)** and **(K-K’’’)** are boxed areas in **(H)** and **(J)**, respectively. DAPI (blue), PatJ (grey) marks the NE (dash lines), Dpn (magenta), GFP (green). Arrowheads indicate NBs. Scale bars: 50 and 10 μm. **L.** Schematic depicting temporal series in the NE. **M-P’’’.** Single section of superficial layer **(M-M’’’, O-O’’’)** WT and **(N-N’’’, P-P’’’)** with *lola^E76^* clones (dash lines) at 72h ALH. GFP (green), DE-Cad (red) marks the NE. **(M-N’’’)** Imp (grey), Mamo (blue). **(O-P’’’)** Syp (grey), Chinmo (blue). Scale bars: 10 μm. **Q.** Corrected Total Cell Fluorescence (CTCF) of Imp and Syp in WT and *lola^E76^* clones at 72h ALH. n=8, 8, 8, and 8 clones. **R.** Summary of *lola* loss-of-function phenotypes in the larval NE compared to WT and *nerfin-1* loss-of-function. Data information: Data are represented as mean ± SEM. *p*-values were obtained using Mann-Whitney test and Wilcoxon matched-pairs signed rank test. \*\*\*\**p* < 0.0001, \*\*\**p* < 0.001, \*\**p* < 0.005, \**p* < 0.05

As tumours generated via the loss-of-function of *nerfin-1* and *lola* caused distinct outcomes in the OL, we hypothesized that Nerfin-1 and Lola regulate different sets of target genes during medulla neurogenesis. To test this hypothesis, we first characterised the expression pattern of Nerfin-1 and Lola using their respective GFP reporters in the OL at wandering L3 stage. Upon NE-NB transition, each NB divides asymmetrically to self-renew and generate GMCs, which in turn divide once to produce medulla neurons (28, 29) (Fig. 4G). The birth order of neurogenesis dictates that the latest born neurons occupying the most superficial position close to NBs and GMCs, and the earliest born neurons occupying the deepest layer. We stained the larval brains for PatJ to mark the apical membrane of NE cells and Dpn to mark NBs. For Lola, we used a GFP against Lola isoform I (Lola.I::GFP) which was previously shown to be expressed in the NBs and neurons of the larval OLs (35). We observed that Lola.I::GFP was expressed in the nuclei of PatJ^+^ NE cells (Fig. 4H-I’’’, yellow dash lines) and Dpn^+^ NBs (Fig. 4H-I’’’, arrowheads). Conversely, Nerfin-1::GFP was not expressed in the NE and the majority of the NBs (Fig. 4J-K’’’). Together, these data suggest that Lola and Nerfin-1 are likely required to function in different cell types in the OL.

To explore target genes downstream of Lola and Nerfin-1, we performed RNA-sequencing and compared the transcriptional profiles of *lola^E76^*, *nerfin-1^159^*, and WT clones from the OLs of wandering L3 larva (Fig. S3D). Our differential expression analyses (adjusted p-value<0.05; fold-change, FC>1.5) revealed 94 differentially expressed genes, including 65 upregulated and 29 downregulated genes in *lola^E76^* compared to WT clones. Between *nerfin-1^159^* and WT clones, there were 164 differentially expressed genes, including 80 upregulated and 82 downregulated genes. Notably, only 11 genes were shared between *lola^E76^* versus WT and *nerfin-1^159^* versus WT comparisons (Fig. S3E). Intriguingly, between *lola^E76^* and WT clones, gene ontology (GO) enrichment for upregulated genes included terms involved in the digestive tract and muscle development (Fig. S3F). These unexpected results suggest that Lola may play a role in suppressing non-CNS genes to ensure the faithfulness of neural identity. Whereas GO enrichment for downregulated genes included synaptic transmission and membrane potential (Fig. S3F), indicating that Lola promotes genes required for neuronal maturation. GO terms enriched for genes upregulated in *nerfin-1^159^*clones included NB division and proliferation (Fig. S3G). Conversely, GO terms enriched for genes downregulated included terms involved in regulation of synaptic transmission and membrane potential (Fig. S3G). Therefore, Nerfin-1 appears to govern neuronal cell fate maintenance by inhibiting self-renewal and proliferation genes while promoting neuronal maturation. Collectively, our data suggest that Lola and Nerfin-1 have different expression patterns and target different sets of genes with minimal overlap in the developing medulla.

### Lola but not Nerfin-1 promotes timely progression of the temporal series in the NE

Since Lola is expressed in the NE (Fig. 4H-I’’’) (35), we next investigated its function in this cell population. Our RNA-seq data showed that *Imp*, *mamo* and *Eip93F* expression levels were significantly altered in *lola^E76^* compared to WT clones (Fig. S3H). In particular, *Imp* was upregulated (FC=79.5) whilst *mamo* and *Eip93F* were downregulated (FC=2.73 and 1.98, respectively). NE cells express a series of temporal factors consisted of Imp, Chinmo, Mamo, Syp, Br and Eip93F that are then inherited by their progenies (16) (Fig. 4L). Imp and Chinmo mark the early temporal window whilst Syp, Br, and Eip93F mark the late temporal window. In addition to our RNA-seq data, we performed immunostaining to examine the protein levels of Imp, Chinmo, Mamo and Syp in the medulla NE cells marked by DE-Cadherin (DE-Cad). At 72 hrs ALH (mid-L3), NE cells in WT clones expressed similar levels of Imp, Chinmo, Mamo, and Syp compared to their counterparts outside of the clones (Fig. 4M-M’’’, O-O’’’). In contrast, *lola^E76^* NE cells exhibited an upregulation of Imp and Chinmo as well as a downregulation of Mamo and Syp compared to NE cells outside of the clones (Fig. 4N-N’’’, P-P’’’). Moreover, quantifications showed that Imp intensity in *lola^E76^*NE cells was significantly higher compared to surrounding WT NE cells (internal control) and vice versa for Syp intensity (Fig. 4Q). Thus, our results suggest that the early temporal window in the NE was extended at the expense of the late temporal window upon Lola inactivation. Intriguingly, we also occasionally found that some *lola^E76^* clones displayed a medially shifted NE/NB border at wandering L3 stage (Fig. S3I-I’, arrowhead), indicative of a delayed NE-NB transition. As opposed to *lola^E76^* clones, the expression of Imp, Chinmo, Syp, and Mamo were not altered in *nerfin-1^159^* NE cells compared to surrounding WT counterparts (Fig. S3J-K’’’). Altogether, our results suggest that only Lola, but not Nerfin-1 is required for the progression of the temporal series in the NE (Fig. 4R).

### Chinmo and Imp promote the persistence of *lola^E76^* dedifferentiated NBs in the pupal OL

As Chinmo is the key determinant that sustains dedifferentiated NB proliferation during pupal life in *nerfin-1^159^*clones, we next examined Chinmo expression in *lola^E76^* clones at wandering L3 stage in the OL. We found that *lola^E76^* clones expressed high levels of Chinmo (Fig. 5A-A’’). To assess whether Chinmo upregulation is functionally required to promote the persistence of *lola^E76^* dedifferentiated NBs in the pupal OL, we knocked down Chinmo in *lola^E76^* clones by RNAi. This manipulation sufficiently reduced Chinmo expression without significantly altering dedifferentiation in mutant clones at wandering L3 stage (Fig. 5A-C). At 48 hrs APF, Chinmo knockdown caused a significant reduction in NB persistence in the OL, and a dispersion of mutant cells, an indicator of neuronal differentiation (24) (Fig. 5D-E’, G). In addition to Chinmo, the RNA-binding protein Imp and Lin-28 are other components of the oncogenic module that defines the oncogenic competency of dedifferentiated NBs in the VNC (26). In our RNA-seq data, we found that differentially upregulated genes in *lola^E76^* clones included both *Imp* and *lin-28* with FC of 79.5 and 2.41, respectively. Moreover, Imp knockdown in *lola^E76^*clones was able to significantly reduce NB persistence at 48 hrs APF, similar to Chinmo knockdown (Fig. 5D-G). Hence, Chinmo and Imp are required to sustain *lola^E76^* dedifferentiated NBs in the pupal OL.

**Figure 5:**
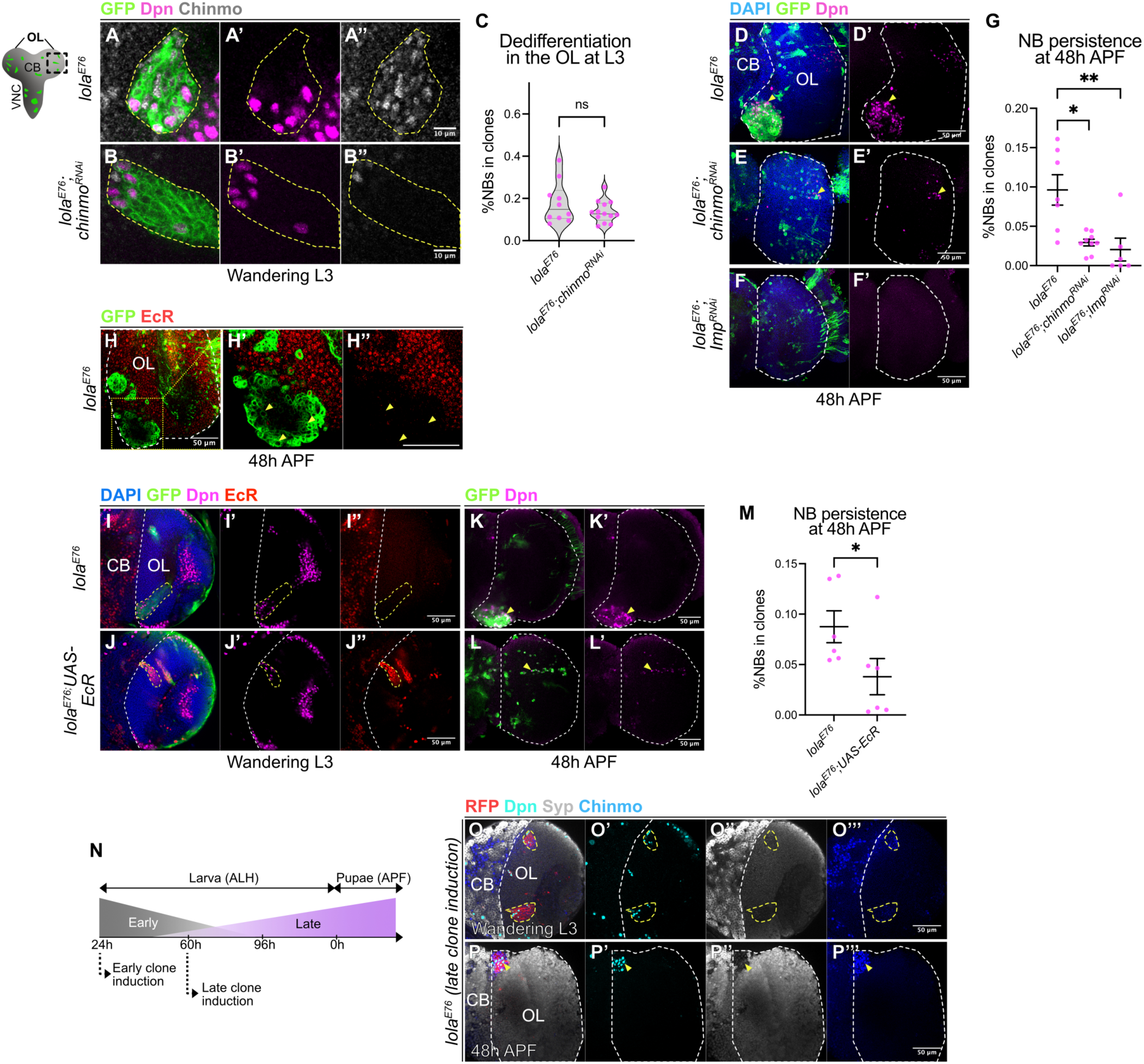
Chinmo and Imp promote the persistence of *lola^E76^* NBs in the pupal OL. **A-B’’.** Single sections of **(A-A’’)** *lola^E76^* and **(B-B’’)** *lola^E76^;chinmo^RNAi^* clones (dash lines) in the OLs (deep layer) at late L3. GFP (green), Dpn (magenta), Chinmo (grey). Scale bars: 10 μm. **C.** NB percentages in *lola^E76^* and *lola^E76^;chinmo^RNAi^* clones in the OLs at late L3. *lola^E76^*: n=10 and 13 clones **D-F’.** Maximum projections of the OLs (dash lines) at 48h APF with **(D)** *lola^E76^*, **(E)** *lola^E76^*;*chinmo^RNAi^* and **(F)** *lola^E76^;Imp^RNAi^* clones. Ectopic NBs (arrowheads). DAPI (blue), GFP (green), Dpn (magenta). Scale bars: 50 μm. **G.** NB percentages in *lola^E76^*, *lola^E76^*;*chinmo^RNAi^,* and *lola^E76^;Imp^RNAi^* clones in the OLs at 48h APF. n=7, 9 and 6. **H-H’’. (H)** Single section of the OL (dash lines) with *lola^E76^* clones at 48h APF. **(H’-H’’)** Boxed areas in (H). Arrowheads indicate GFP^+^ cells. GFP (green), EcR (red). Scale bars: 50 μm. **I-J’’.** Single section in the deep layers of the brain lobes at late L3 with **(I-I’’)** *lola^E76^*, and **(J-J’’)** *lola^E76^*;*UAS-EcR* clones (yellow dash lines). DAPI (blue), GFP (green), Dpn (magenta), EcR (red). White dash lines indicate the CB/OL border. Scale bars: 50 μm. **K-L’.** Single section of the OLs (dash lines) at 48h APF with **(K-K’)** *lola^E76^*, and **(L-L’)** *lola^E76^*;*UAS-EcR* clones (arrowheads). GFP (green), Dpn (magenta). Scale bars: 50 μm. **M.** NB percentages in *lola^E76^* and *lola^E76^*;*UAS-EcR* clones in the OLs at 48h APF. n=6 and 6. **N.** Clonal induction schematic. **O-O’’’.** Single section of the brain lobe (deep layer) with *lola^E76^* clones (yellow dash lines) at late L3. RFP (red), Dpn (cyan), Syp (grey), Chinmo (blue). White dash lines indicate the CB/OL border. Scale bars: 50 μm. **P-P’’’.** Maximum projection of OL (dash lines) with *lola^E76^* clones (arrowheads) at 48h APF. RFP (red), Dpn (cyan), Syp (grey), Chinmo (blue). Scale bars: 50 μm. Data information: Data are represented as mean ± SEM. *p*-values were obtained using Mann-Whitney test and Kruskal-Wallis test with Dunn’s test to correct for multiple comparisons. \*\**p* < 0.005, \**p* < 0.05

### Chinmo upregulation occurs independently of ecdysone and the temporal identity of its cell of origin upon Lola inactivation

We next aimed to understand how Chinmo is regulated in *lola^E76^*clones to sustain the proliferation of dedifferentiated NB in the pupal OL. As ecdysone signalling mediates the suppression of Chinmo to eliminate *nerfin-1^159^*dedifferentiated NBs in the pupal OL, we wondered how *lola^E76^* dedifferentiated NBs were able to retain Chinmo expression in spite of their location. Unlike *nerfin-1^159^* clones that express EcR (Fig. 3L-L’’), we found that EcR expression was downregulated in *lola^E76^* clones compared to WT surrounding cells in the OL at 48 hrs APF (Fig. 5H-H’’, arrowheads). As such, it is likely that *lola^E76^* dedifferentiated NBs in the OL are not receptive to ecdysone signals that normally promote Chinmo silencing. To test if ecdysone signalling is sufficient to promote an elimination of *lola^E76^* dedifferentiated NBs in the pupal OL, we overexpressed both isoforms A and B of EcR in *lola^E76^* clones (Fig. 5I-J’’). At 48 hrs APF, while *lola^E76^* clones exhibited persistent ectopic NBs forming tumours in the OL, *lola^E76^;UAS-EcR* clones showed significantly less persistent NBs and the clones exhibited a dispersed morphology (Fig. 5K-M). Therefore, the absence of EcR in *lola^E76^* clones facilitates the persistence of dedifferentiated NBs in the pupal OL.

We have shown that Lola promotes the progression of the early Imp^+^Chinmo^+^ to late Syp^+^ temporal identity in the NE (Fig. 4R). In *lola^E76^* clones, we observed that Chinmo was upregulated in the NBs on the superficial layers of the OL medulla at wandering L3 stage (Fig. S4A-A’’’ yellow arrowheads). This indicates that NBs and their progenies inherit the temporal identity of the NE cells in *lola^E76^*clones. Thus, it is possible that *lola^E76^* dedifferentiated NBs express Chinmo as they are generated during the Chinmo^+^ time window (24 hrs ALH, early time point, Fig. 5N). To test this hypothesis, we induced *lola^E76^* clones at 60 hrs ALH (late time point, Fig. 5N) to bypass the Chinmo^+^ time window. Despite being generated in the Chinmo^-^ time window, *lola^E76^* clones continued to express the early temporal marker Chinmo and failed to turn on the late temporal marker Syp at wandering L3 stage (Fig. 5O-O’’’). At 48 hrs APF, *lola^E76^* ectopic NBs were persistent in the OL (Fig. 5P-P’’’). Thus, our data suggest that *lola^E76^* dedifferentiated NBs can persist independently of whether they are generated in the Chinmo^+^ or Chinmo^-^ temporal window.

Furthermore, we attempted to test if inhibiting the early temporal identity in the NE can prevent the persistence of *lola^E76^* dedifferentiated NBs in the pupal OL. To do so, we overexpressed Syp in *lola^E76^* clones, which was shown to repress the expression of Chinmo and Imp in the CB/VNC (Liu et al., 2015; Ren et al., 2017; Yang et al., 2017). However, at 48 hrs APF, Syp overexpression was neither sufficient to downregulate Imp, nor able to suppress the persistence of *lola^E76^* ectopic NBs in the OL (Fig. S4B-D). Thus, it is possible that the regulatory relationship between Imp-Chinmo-Syp in the OL may differ from that of the CB/VNC.

### Notch signalling is not required for neuronal dedifferentiation in *lola^E76^* clones

We and others previously showed that neuronal dedifferentiation induced via *nerfin-1* loss-of-function in the OL requires Notch signalling activation (19, 22). We wondered if Notch signalling is also required for neuronal dedifferentiation induced by *lola* loss-of-function. Our RNA-seq data showed that while *Notch* (*N*) was upregulated (FC=2.08) in *lola^E76^* compared to WT cells, *Delta* which encodes for a Notch ligand, as well as the downstream effector of Notch signalling, *E(spl)mγ* were not significantly altered (Fig. S4E). In line with this, immunostaining in the OL showed that the intracellular domain of Notch (NICD) was upregulated in *lola^E76^* clones whereas Delta appeared not altered compared to WT cells surrounding the clones at the wandering L3 stage (Fig. S4F-G’’). To address if Notch signalling is necessary for neuronal dedifferentiation induced by *lola* loss-of-function, we knocked down *N* via RNAi in *lola^E76^* clones. This manipulation did not significantly alter the rate of dedifferentiation at wandering L3 stage compared to the *lola^E76^* control (Fig. S4H-J). Together, our data suggest that Notch signalling does not mediate neuronal dedifferentiation induced via Lola inactivation.

### *lola^E76^* dedifferentiated NBs exhibited delayed temporal progression in the OL

Medulla NBs sequentially express a series of tTFs including Homothorax (Hth), Eyeless (Ey), Sloppy paired (Slp), Dichaete (D), and Tailless (Tll) as they age (36) (Fig. 6A). We next aimed to investigate whether this temporal series is altered in *lola^E76^* dedifferentiated NBs using three representative tTFs: the early-tTF Ey, mid-tTF Slp and late-tTF Tll. It was previously shown that *lola* knockdown in medulla NBs caused delayed temporal progression during late larval stages (35). Consistent with this, at wandering L3 stage, we observed that Ey, Slp, and Tll expression in *lola^E76^* NBs appeared reduced compared to WT NBs on the superficial layers, indicative of delayed temporal progression (Fig. 6B-F’’’). Thus, our data confirms the role of Lola in controlling the speed of temporal progression in the medulla NBs. In the deep layers of the larval medulla where WT neurons and dedifferentiated NBs are located, we observed that a subset of cells in *lola^E76^* clones expressed Ey and Slp, but not Tll at wandering L3 stage (Fig. 6B, G-H’’’). By 48 hrs APF, fewer dedifferentiated NBs expressed Ey and Slp, and more cells expressed Tll in *lola^E76^* clones (Fig. 6I-K). Together, our data suggest that *lola^E76^* dedifferentiated NBs exhibited delayed progression through the temporal series (Fig. 6L).

**Figure 6:**
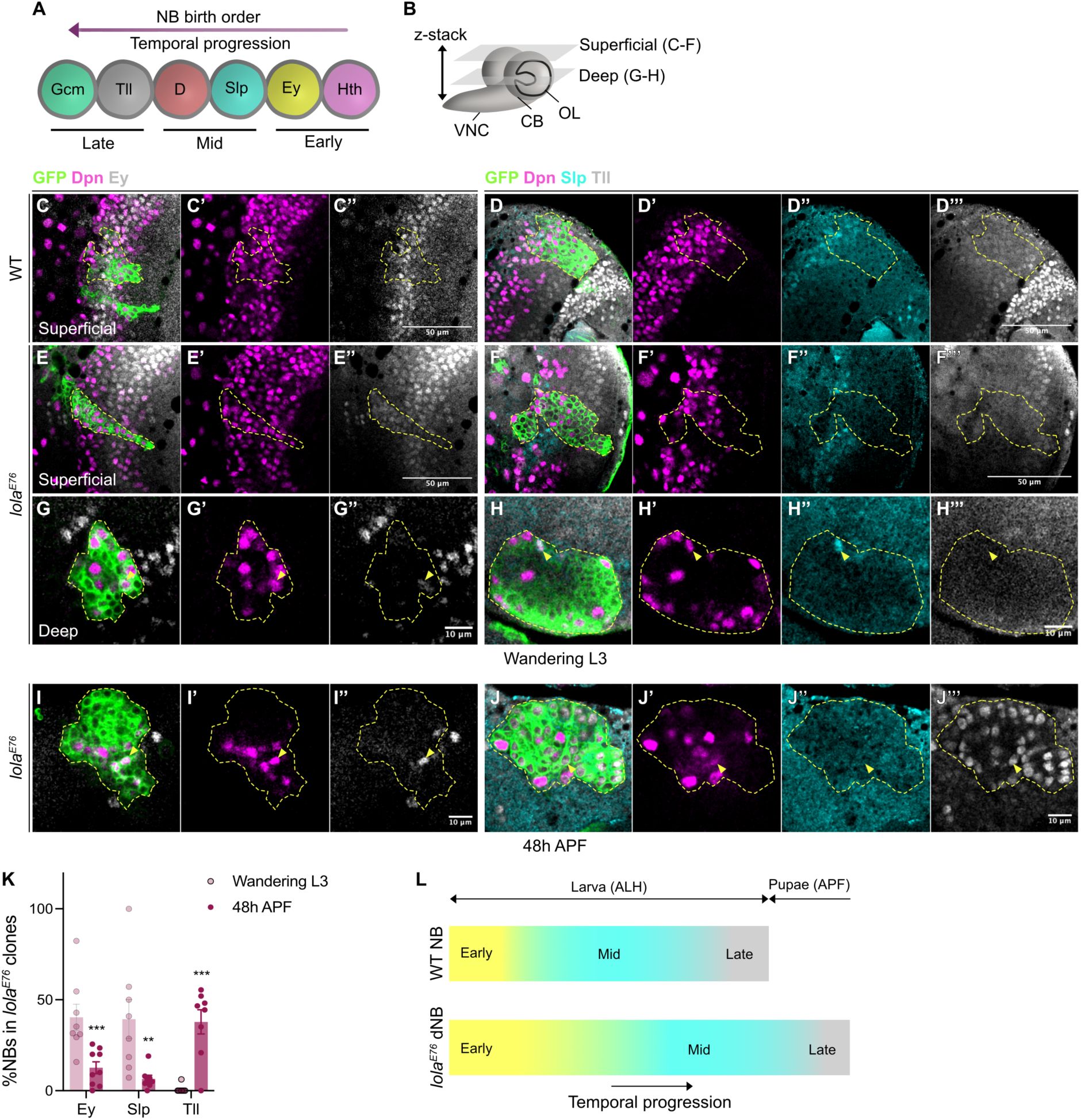
Temporal progression is delayed in *lola^E76^* dedifferentiated NBs. **A.** Schematic depicting the temporal series in the medulla NBs. **B.** Schematic depicting the side view of the larval CNS. Representative confocal images of the larval brain are taken from either superficial (for Fig. 6C-F) or deep (for Fig. 6G-H) layers. **C-F’’’.** Single confocal sections of **(C-D’’’)** WT and (**E-F’’’**) *lola^E76^* clones (dash lines) on the superficial layers of the medulla at wandering L3 stage. **(C-C’’, E-E’’)** GFP (green), Dpn (magenta), Ey (grey). **(D-D’’’, F-F’’’)** GFP (green), Dpn (magenta), Slp (cyan), Tll (grey). Scale bars: 50 μm. **G-H’’’.** Single confocal sections of *lola^E76^* clones (dash lines) in the deep layers of the medulla at wandering L3 stage. **(G-G’’)** GFP (green), Dpn (magenta), Ey (grey). Arrowheads indicate NBs expressing Ey. **(H-H’’’)** GFP (green), Dpn (magenta), Slp (cyan), Tll (grey). Arrowheads indicate NBs expressing Slp. Scale bars: 10 μm. **I-J’’’.** Single confocal section of *lola^E76^* clones (dash lines) in the OL at 48h APF. **(I-I’’)** GFP (green), Dpn (magenta), Ey (grey). Arrowheads indicate NBs expressing Ey. **(J-J’’’)** GFP (green), Dpn (magenta), Slp (cyan), Tll (grey). Arrowheads indicate NBs expressing Tll. Scale bars: 10 μm. **K.** Quantifications of the NB ratio expressing Ey, Slp, and Tll in *lola^E76^*clones at wandering L3 stage and 48h APF in the OLs. n=8, 9, 8, 8, 8, and 8 clones. **L.** Summary schematic depicting the delayed temporal progression in *lola^E76^*dedifferentiated NBs (dNBs) compared to WT NBs. Data information: Data are represented as mean ± SEM. *p*-values were obtained using Multiple Mann-Whitney test and Kruskal-Wallis test with Dunn’s test to correct for multiple comparisons. \*\*\**p* < 0.001, \*\**p* < 0.005.

To test the requirement of tTF expression for *lola*-mediated dedifferentiation and tumorigenesis in the OL, we manipulated three tTFs - Hth, Ey, and Slp in *lola^E76^* clones. Upon the overexpression of Hth and inhibition of Ey using validated fly lines (35, 37), we found that the rate of dedifferentiation was not significantly altered compared to *lola^E76^* control clones at wandering L3 stage (Fig. S5A-C’, E). We also knocked down Slp by *slp1^RNAi^* (knockdown efficiency validated in fig. S5F-F’’) in *lola^E76^* clones. Similar to Hth overexpression and Ey knockdown, Slp knockdown did not significantly alter the proportion of NBs in *lola^E76^* clones (Fig. S5D-E). Accordingly, our data suggest that the expression of these tTFs is dispensable for the modulation of neuronal dedifferentiation caused by Lola inactivation. At 48 hrs APF, *lola^E76^* clones exhibited persistent NBs (Fig. S5G-G’) and this phenotype was not significantly altered by the manipulations of Hth, Ey and Slp (Fig. S5H-K). Taken together, our data suggest that although *lola^E76^* NBs display a delayed temporal progression from larval to pupal development, tTFs play minor roles in promoting NB persistence and tumorigenesis in the OL.

### Lola expression in *nerfin-1* loss-of-function clones promotes the elimination of dedifferentiated NBs

Given that both Nerfin-1 and Lola are required to maintain the differentiated state of medulla neurons, we wondered if these factors acted redundantly. To test this, we examined the expression of Lola.I::GFP reporter in *nerfin-1^RNAi^* flip-out clones at wandering L3 stage by immunostaining. We showed that Lola expression was not altered in *nerfin-1^RNAi^* clones compared to surrounding WT cells in the medulla (Fig. 7A-A’’’). Similarly, *lola* mRNA expression level was comparable between *nerfin-1^159^* and WT clones (Fig. 7B). To test if Lola is functionally required in *nerfin-1^159^*dedifferentiated NBs, we knocked down Lola in *nerfin-1^159^* clones by RNAi. In *nerfin-1^159^, lola^RNAi^,* and *lola^RNAi^;nerfin-1^159^* clones, we observed ectopic NBs in the deep layers of the medulla at wandering L3 stage (Fig. 1C, 7C-D’), indicative of neuronal dedifferentiation. By 48 hrs APF, *lola^RNAi^ and lola^RNAi^;nerfin-1^159^* clones, but not *nerfin-1^159^* clones displayed persistent ectopic NBs in the OLs (Fig. 1M-M’, 7E-F’). Notably, quantifications showed that NB persistence in *lola^RNAi^;nerfin-1^159^*clones was significantly higher than that of *lola^RNAi^* clones at 48 hrs APF (Fig. 7G).

**Figure 7:**
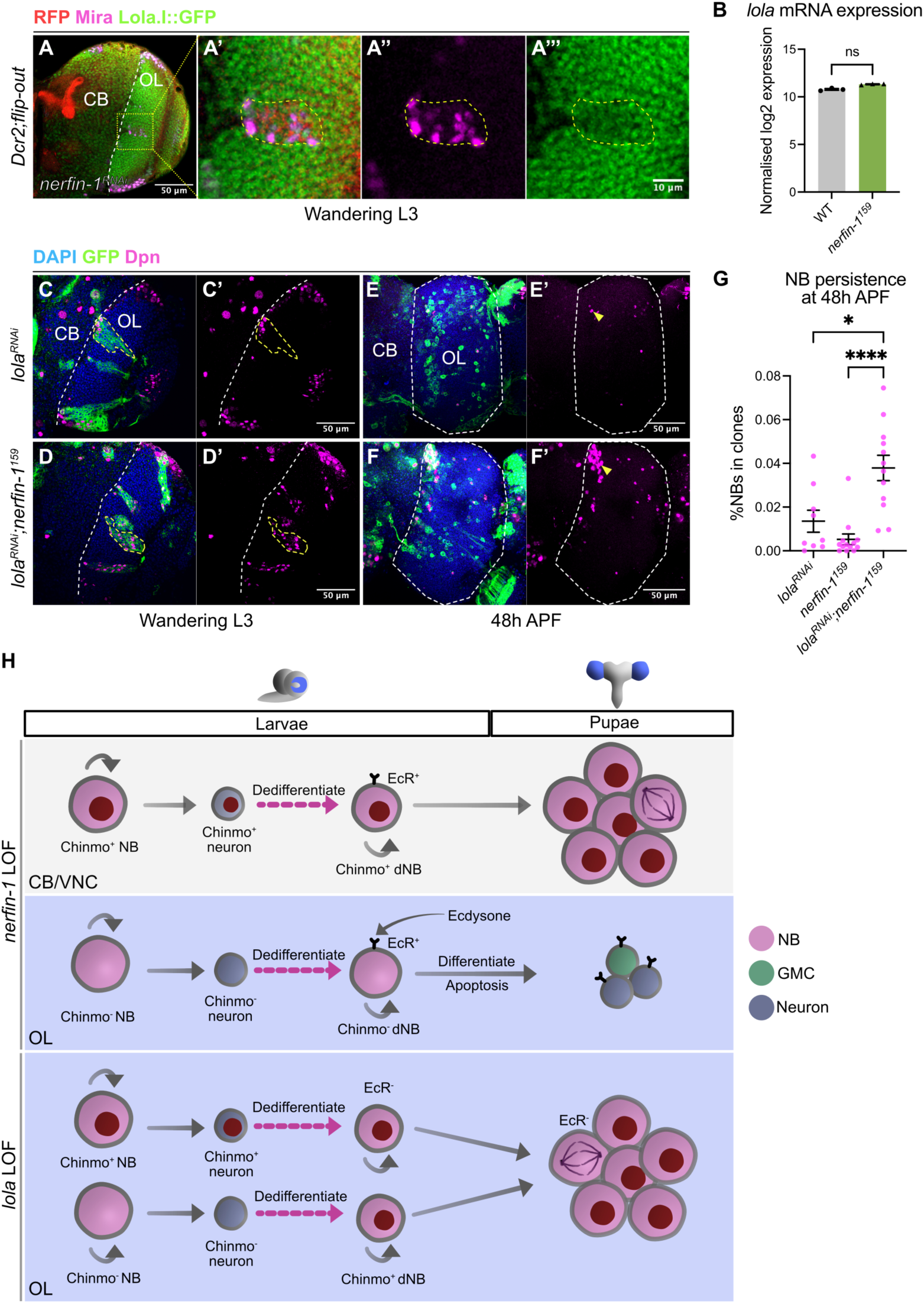
Lola expression in *nerfin-1* loss-of-function clones promotes elimination of ectopic NBs in the pupal OL. **A-A’’’. (A)** Single confocal section in the deep layers of the brain lobe at wandering L3 stage with *nerfin-1^RNAi^* flip-out clones. Scale bars: 50 μm. **(A’-A’’’)** is boxed area in (A). RFP (red), Mira (magenta). Scale bars: 10 μm. Lola.I::GFP (green). Yellow dash lines indicate mutant clones. White dash lines indicate the CB/OL borders. **B.** *lola* normalised log2 expression from RNA-seq data of WT and *nerfin-1^159^*cells collected from the late larval OLs. n=3 and 3 samples. **C-D’.** Single confocal sections in the deep layers of the brain lobes at wandering L3 stage with **(C-C’)** *lola^RNAi^* and **(D-D’)** *lola^RNAi^*;*nerfin-1^159^* clones (yellow dash lines). DAPI (blue), GFP (green), Dpn (magenta). White dash lines indicate the CB/OL borders. Scale bars: 50 μm. **E-F’.** Maximum projections of the OLs (dash lines) at 48h APF with **(E-E’)** *lola^RNAi^* and **(F-F’)** *lola^RNAi^*;*nerfin-1^159^* clones. DAPI (blue), GFP (green), Dpn (magenta). Arrowheads indicate ectopic NBs in the OLs. Scale bars: 50 μm. **G.** Quantifications of the NB percentages of *nerfin-1^159^*, *lola^RNAi^* and *lola^RNAi^;nerfin-1^159^* clones in the OLs at 48h APF. n=13, 9 and 12. **H.** Summary schematic: In the CB/VNC, Chinmo expression inherited from dedifferentiating *nerfin-1* loss-of-function (LOF) neurons promotes persistence of dedifferentiated NBs (dNBs) during pupal stages. In the OL, ecdysone signalling represses *chinmo* expression in *nerfin-*1 LOF clones, resulting in the elimination of dNBs through terminal differentiation and apoptosis. *lola* LOF clones in the OL ectopically upregulates Chinmo. Due to the absence of EcR in *lola* LOF clones, Chinmo expression is not suppressed, resulting in dNB persistence in the pupal OL. Data information: Data are represented as mean ± SEM. *p*-values were obtained using Mann-Whitney test and Kruskal-Wallis test with Dunn’s test to correct for multiple comparisons. \*\*\**p* < 0.001, \**p* < 0.05.

In the VNC, *lola^RNAi^;nerfin-1^159^* clones exhibited no significant difference in the percentages of NBs at both wandering L3 stage and at 48 hrs APF (Fig. S6A-F), indicating that Lola is dispensable for *nerfin-1* loss-of-function tumour overgrowth in this region. Collectively, our results suggest that Lola promotes the elimination of dedifferentiated NBs in *nerfin-1* loss-of-function clones in the OL but not the CB/VNC during pupal development.

## Discussion

Dedifferentiation of mature specialised cells into stem-progenitor cells that can self-renew and re-differentiate provides valuable opportunities for regeneration. However, studies have shown that dedifferentiation can produce cancer stem cells that drive malignant tumour overgrowth and heterogeneity (38). Thus, it is crucial to understand what factors determine these contrasting outcomes of dedifferentiation. In this work, we have discovered that oncogenic tTFs are key determinants of this process. In the CB/VNC, *nerfin-1^159^*dedifferentiated NBs express the oncogenic tTF Chinmo that promotes their persistence during pupal development. Conversely, in the OL, active ecdysone signalling promotes *chinmo* silencing. The absence of Chinmo in turn enables the elimination of *nerfin-1^159^* dedifferentiated NBs by apoptosis and terminal differentiation. *lola^E76^* dedifferentiated NBs on the other hand are not responsive to ecdysone signals during pupal development, thereby ectopically express Chinmo and persist in the pupal OL. Together, these mechanisms dictate the outcome of neuronal dedifferentiation (Fig. 7H). Our study offers evidence that genetic mutations as well as region-specific cues can contribute to the oncogenic competence of stem-progenitor cells derived from dedifferentiation.

### Chinmo is the key determinant of oncogenic competence in the CNS

Our data suggest that the differential expression and regulatory mechanism of Chinmo can dictate the oncogenic outcome of dedifferentiated NBs; such that high Chinmo induce oncogenesis whilst no/low Chinmo allows elimination of the dedifferentiated progenitor pool. An outstanding question that remains to be addressed is the mode-of-action of Chinmo. In the WT CB/VNC, Chinmo acts as a tTF that defines the early temporal window in the NBs and their progenies, but not a regulator of NB proliferation (11, 26). In *pros^-/-^*dedifferentiated tumours in the VNC, Chinmo was suggested to promote cell division and protein synthesis (26). However, in the OL, Chinmo does not only act as a tTF in the NE progenitors and its progeny but also governs NE cell self-renewal, and thereby, the timing of NE-NB transition (17). Therefore, whether and how the function of Chinmo differs between the CB/VNC and the OL need further investigation.

### Steroid hormone signalling promotes elimination of dedifferentiated NBs by suppressing oncogenic tTFs

In the developing OL medulla, Chinmo expression is silenced in the NE by a pulse of ecdysone at critical weight which coincides with the acceleration of the NE-NB transition, followed by post-embryonic neurogenesis (17). As such, most medulla neurons are generated post-critical weight, during the Chinmo^-^time window and are thus incompetent to initiate tumour overgrowth following dedifferentiation. However, because *nerfin-1^-/-^* clones were induced during the early Chinmo^+^ temporal window of the medulla, it was unexpected that *nerfin-1^-/-^* dedifferentiated NBs did not inherit Chinmo expression. Our data showed that *nerfin-1^-/-^* clones expressed EcR and thereby, are responsive to ecdysone signals between larval-pupal development. This in turn promotes *chinmo* silencing, resulting in the elimination of dedifferentiated NBs. Nevertheless, it remains to be addressed whether ecdysone signalling necessitates either or both the initiation and the maintenance of Chinmo silencing in this case. In addition, how ecdysone signalling controls Chinmo expression in the OL is yet to be interrogated. In the developing medulla, ecdysone signalling at the mid-larval stage induces expression of its target gene *br* in the NE to promote the NE-NB transition (18). In the wing imaginal disc, ecdysone signalling induces a Chinmo-to-Br expression switch at the late larval stage (39). In addition, Br and Chinmo inhibit each other’s expression (39, 40). As such, it is possible that like the wing disc, ecdysone signalling inhibits Chinmo expression in the *nerfin-1^-/-^* clones via its downstream effector Br. In *lola^-/-^* clones, despite their location in the OL, dedifferentiated NBs continued to divide during late pupal stages (Fig. 4E-F). Unlike *nerfin-1^-/-^* clones, we found that *lola^-/-^* clones did not express EcR and EcR misexpression is sufficient to eradicate *lola^-/-^*tumorigenesis in the pupal OL (Fig. 5H-M). Nevertheless, further investigation is needed to better understand how Lola regulates EcR expression. In the developing CB/VNC, it was previously shown that ecdysone signalling is dispensable for Chinmo expression in the NBs (17), which may account for the persistence of the *nerfin-1^-/-^*dedifferentiated NBs in these regions.

### Lola regulates temporal progression in the medulla NE and dedifferentiated NBs

We and other labs have shown that two Lola isoforms Lola.I and Lola.T are expressed in the larval NE (35) (Fig. 4H-I’’’). Here, our work has demonstrated that Lola regulates the progression of the early-to-late temporal window in the NE progenitor pool. Intriguingly, some *lola^-/-^*clones exhibited a delayed NE-NB transition (Fig. S3I-I’). Given that the expression of tTFs has been implicated in the regulation of NE self-renewal and differentiation (17, 18), it is likely that Lola also governs the timing of the NE-NB transition in the larval OL. As such, the previously demonstrated delay in temporal progression in the medulla NBs upon Lola inactivation (35) might be a consequence of a delay in the NE-NB transition. Additionally, we report that *lola^-/-^* dedifferentiated NBs exhibit delayed temporal progression (Fig. 6G-L). While it has been suggested that the competence to generate brain tumours is determined by temporal identity (26, 41), our results suggest that temporal progression of *lola^-/-^* dedifferentiated NBs plays a minor role in their persistence and tumorigenicity (Fig. S5).

It is of note that we used *lola^E76^* null mutation which removes all 12 Lola isoforms (34). This left us unable to identify which Lola isoform functions in the NE, or to dissect the specific roles of each Lola isoforms in different cell types (NE, NBs, neurons) in the medulla. Previous work showed that Lola-N overexpression can rescue tumour overgrowth caused by *lola^E76^* dedifferentiated NBs in the adult OL (20). However, Lola-N mutant clones displayed no ectopic NBs in the adult OL (42), suggesting that there is functional redundancy between Lola isoforms that needs to be taken into consideration.

### Differential target genes of Nerfin-1 and Lola in the developing medulla

Our RNA-seq analyses have revealed that there is very little overlap between differential expressed genes in *nerfin-1^-/-^* and *lola^-/-^* clones compared to the WT (Fig. S3E). This may account for the contrasting tumorigenic phenotype between two mutations in the OL. The differentially expressed genes are enriched in partially overlapping GO terms, in both cases, GO enrichment for downregulated genes involves neuronal function and maturation (Fig. S3F-G). In *nerfin-1^-/-^*, GO enrichment for upregulated genes includes NB division and proliferation (Fig. S3G). This is expected given the expression pattern of Nerfin-1 in immature neurons in the developing medulla (19, 22). Interestingly, GO enrichment for upregulated genes in *lola^-/-^* clones are enriched in non-CNS genes (Fig. S3F). This echoes recent findings regarding the chromatin remodeller Mi-2 and the condensin I subunit Cap-G whose knockdowns in neurons cause ectopic upregulation of genes signatures of the fat body, the hindgut, or cilium assembly, etc, that tend to not associate with the CNS (43, 44). Accordingly, it will be of interest to probe how Lola represses the expression of organ and lineage inappropriate genes in the neurons, either via chromatin remodelling or other mechanisms. A caveat in our study lies in transcriptomic profiling using bulk RNA-seq. As we have demonstrated that Nerfin-1 and Lola are expressed and function differently in different cell types of the developing medulla, it is unlikely that bulk RNA-seq precisely captured the gene network dynamic as well as the heterogeneity of *nerfin-1^-/-^*and *lola^-/-^* clones.

Characterisations of paediatric brain tumours such as glioblastoma and medulloblastoma have revealed the link between the oncogenic mutations, the tumour location and the overall prognosis in patients (5–7). In glioblastoma, two mutations of the *H3F3* gene that lead to two amino acid substitutions of the histone 3 variant 3, H3.3-G34R and H3.3-K27M induce tumour overgrowth in anatomical distinct regions of the CNS. While the H3.3-G34R subtype is mostly found in the brain hemisphere, the H3.3-K27M subtype is often found in the hindbrain (6, 7). A recent study using compartmentalised NSC culture derived from anatomical relevant areas showed that high expression of the transcription factor FOXG1 in the forebrain confers its housing NSCs the competency to undergo H3.3-G34R-induced oncogenic transformation (45). Hence, differential gene expression profiles between brain regions appear to underpin their varying susceptibility to oncogenic transformation. Our investigation further suggests that elucidating the regionalised control of temporal patterning can aid us in understanding the outcome of dedifferentiated tumours; however, the result needs to be considered within the context of a specific mutation.

## Materials and Methods

### Fly stocks and husbandry

The fly strains utilised are detailed Table 1.

**Table 1:** Fly stock resources table.

| System | Genotype | Source |
| --- | --- | --- |
| Flip-out | *hs-FLP, act>CD2>GAL4; UAS-Dcr2, UAS-GFP* | Lei Zhang |
|  | *hsFLP; actc5>CD2>GAL4, UAS-GFP/TM6B* | (21) |
| MARCM | *yw, hs-FLP; tub-GAL4, UAS-mCD8::GFP; FRT2A, tub-GAL80* | Alex Gould |
|  | *UAS-mCD8::GFP, hsFLP; FRT42D, tub-GAL80; tub-GAL4* | Helena Richardson |
|  | *hs-FLP; FRT42D, tub-GAL80; tub-GAL4, UAS-mCD8::RFP* | Kieran Harvey |
|  | *FRT42D, lola<sup>E76</sup>* | Tony Southall |
|  | *Df(3L)nerfin-1<sup>159</sup>* | Ward Odenwald |
|  | *FRT2A* | Alex Gould |
| GAL4/GAL80 | *ey<sup>R16F10</sup>-GAL4* | BL48737 |
|  | *tub-GAL80<sup>ts</sup>* | BL7108 |
| Reporter | *Nerfin-1::GFP* | BL67385 |
|  | *Lola.l::GFP* | BL38662 |
|  | *UAS-mCherry-chinmo<sup>UTRs</sup>* | Cedric Maurange |
|  | *chinmo-lacZ* | BL10440 |
| RNAi | *UAS-pros RNAi* | BL42538 |
|  | *UAS-gcm RNAi* | VDRC110539 |
|  | *UAS-chinmo RNAi (III)* | BL33638 |
|  | *UAS-chinmo RNAi (II)* | This study |
|  | *UAS-EcR<sup>DN</sup>* | BL9451 |
|  | *UAS-Imp RNAi* | BL34977 |
|  | *UAS-nerfin-1 RNAi* | VDRC101631 |
|  | *UAS-lola RNAi* | VDRC101925 |
|  | *UAS-lola RNAi* | BL26721 |
|  | *UAS-mCherry RNAi* | BL35784 |
|  | *UAS-lacZ RNAi* | Kieran Harvey |
|  | *UAS-ey RNAi* | BL32486 |
|  | *UAS-slp1 RNAi* | BL29354 |
| Overexpression | *UAS-p35* | BL5072 |
|  | *UAS-Myc* | BL9674 |
|  | *UAS-chinmo<sup>FL</sup>* | BL50740 |
|  | *UAS-syp<sup>RB</sup>::HA* | Tzumin Lee |
|  | *UAS-Imp.G* | BL93386 |
|  | *UAS-mCherry-let7<sup>Sponge</sup>* | BL61365 |
|  | *UAS-EcR A+B* | Leonie Quinn |
|  | *UAS-hth* | BL98446 |
| attP docking site | *P{CaryP}Msp300[attP40] (II)* | BL25709 |

For clonal analysis, flip-out clones were induced at 24 hrs ALH, for 10 min at 37°C then moved to 29°C until dissection. This is except for experiments with *UAS-chinmo^FL^*whereby clones were induced at 48 hrs ALH for 8 min and larva were allowed to develop until dissection at room temperature to prevent animal lethality. Mosaic Analysis with a Repressible Cell Marker (MARCM) (46) clones were induced at 24 hrs ALH unless indicated otherwise, for 30 min at 37°C and then moved to 29°C until dissection. This is except for experiments with *UAS-pros^RNAi^* in which clones were induced for 15 min and animals were kept at 25°C to prevent excessive tumour overgrowth in the CNS.

For temporally restricted GAL4 expression by GAL80^ts^, embryos were collected overnight at room temperature and then moved to restrictive 18°C for 5 days. Larva were next transferred to permissive 29°C to allow GAL4-induced transgene expression and dissected at either wandering L3 stage or 24 hrs APF.

### Generation of *UAS-chinmo^RNAi^* (II) fly stock

To generate *chimmo^RNA^*^i^ fly stock, oligos were designed based on the *UAS-chinmo^RNAi^* TRiP line (BL33638), with sense and anti-sense sequences as 5’-AGCGCGAATATCGTTGACAGA-3’ and 5’-TCTGTCAACGATATTCGCGCT-3’, respectively. After annealing, the resulting DNA fragment was cloned into VALIUM20 vector linearized by NheI and EcoRI. The ligation product was transformed into Stable Competent *E.Coli* (NEB C3040H). Transformed colonies were selected by the presence of pVALIUM20 vector by PCR with forward primer 5’-ACCAGCAACCAAGTAAATCAAC-3’ and reverse primer 5’-TAATCGTGTGTGATGCCTACC-3’. Plasmids were purified from selected colonies using Isolate II Plasmid Miniprep (Meridian Bioscience) for sequencing and injected into flies carrying an attP40 docking site.

### Immunostaining

Larval and pupal brains were dissected in phosphate buffered saline (PBS), fixed in 4% formaldehyde in PBS for 20 min and rinsed three times in 0.5% Triton X-100 in PBS (PBST) at room temperature. For immunostaining, brains were incubated in primary antibodies overnight at 4°C, followed by two 0.5% PBST washes and then an overnight secondary antibody incubation at 4°C. Brains were next washed twice with 0.5% PBST and optically cleared in 50% glycerol for 20 min. Samples were mounted in either 80% glycerol or anti-fade mounting medium containing 4% n-propyl gallate, on glass slides and sealed with coverslips (22×22 mm, No 1.5) for image acquisition. The primary antibodies were: rat anti-Dpn (1:200, Abcam 195172), chick anti-GFP (1:1000, Abcam 13970), rat anti-Elav (1:50, DSHB 7E8A10), mouse anti-Repo (1:50, DSHB 8D12), rabbit anti-Dcp-1 (1:100, Cell Signalling 95785), rabbit anti-Myc (1:100, Santa Cruz Biotech d-1-717), rabbit anti-RFP (1:200, Rockland #600-401-379), mouse anti-Mira (1:100, a gift from Alex Gould), rabbit anti-PatJ (1:500, a gift from Helena Richardson), rat anti-DE-Cad (1:200, DSHB DCAD2), rat anti-Chinmo (a gift from Nicholas Sokol), guinea pig anti-Chinmo & rabbit anti-Syp & guine pig anti-Mamo (1:200, gifts from Claude Desplan), mouse anti-EcR common (EcR^comm^) (1:50, DSHB Ag10.2), rabbit anti-Imp (1:500, a gift from Paul M Macdonald), mouse anti-NICD (1:50, DSHB C179C6), mouse anti-Delta (1:10, DSHB C5949B), mouse anti-Ey (1:50, DSHB), guinea pig anti-Slp & rabbit anti-Tll (1:200, gifts from Kuniaki Saito), Phalloidin 647 (1:10,000 Abcam 176759). Secondary donkey antibodies conjugated to Alexa 555, Alexa 647, goat antibodies conjugated to Alexa 405, 488, 555, and 647 (Molecular Probes and Abcam) were used at 1:500.

### Image acquisition and processing

Images were acquired using Olympus FV3000 confocal microscope with 40X (NA 0.95, UPSALO), and 60X (NA 1.30, UPLSAPO) objectives. Δz=1.5 μm. Images were processed using Fiji (https://imagej.net/Fiji). Quantification was performed in Fiji or via 3D reconstruction in Imaris (Bitplane). To quantify the percentages of NBs in clones at the larval stage or in the pupal OL in Imaris, GFP^+^ volume were first detected and measured. The GFP^+^ volume were then utilised to make a mask for Dpn^+^ cells which was in turn used to measure the number of NB. The percentages of NBs in clones were calculated as the number of GFP^+^Dpn^+^ cells/total GFP^+^ volume x 100%. To quantify the percentages of neuron volume in Imaris, GFP^+^ volume in the whole OL or VNC were first detected and measured. The GFP^+^ volume were then utilised to make a mask for Elav^+^ cells which was in turn used to measure total Elav^+^ volume. The percentages of neuron volume in clones were calculated as GFP^+^Elav^+^/total GFP^+^ volume x 100%. Quantifications of the percentages of NBs expressing tTFs (Ey, Slp, and Tll) in *lola^E76^* clones were done manually in Fiji. Quantifications of the NB number in the OL at 24 hrs APF were done either manually in Fiji or Imaris. Quantifications of Imp and Syp intensity in *lola^E76^* and WT cells around *lola^E76^*clones were done in Fiji using Corrected Total Cell Fluorescence (CTCF) method. CTCF = Integrated Density – (Area of selected cell x Mean fluorescence of background readings). Images were assembled in Affinity Publisher 2. Schematics were created in Affinity Design 2. Scale bars = 10, 20, or 50 μm per indicated.

### RNA-sequencing

*FRT42D* (WT), *FRT2A,nerfin-1^159^*, *FRT42D,lola^E76^*mutant cells were isolated from wandering L3 stage OLs by FACS purification using previously published protocol (47). Briefly, OLs were dissected from the larval CNS in cold Schneider’s culture medium (Gibco) and dissociated using papain and collagenase I (Sigma-Aldrich). Cells were sorted by flow cytometry into lysis buffer and extracted for RNA using Single cell purification kit (Norgen Biotek SKU51800). Samples from sorts executed on different days were pooled into three samples per genotype, containing 33,000-105,000 cells (about 15-70 ng total RNA) per sample. RNA concentration and quality were analysed by Tapestation Bioanalyzer 4150. Libraries were amplified and prepared with QuantSeq (Lexogen) following standard protocol. RNA-sequencing was carried out on NextSeq2000 instrument (Illumina), using 100 base single-end reads and 100 cycles. RNA-seq samples consisted of three replicates per genotype and were all sequenced in the same batch.

### Bioinformatic analyses

Alignment of the RNA-seq data to the *Drosophila melanogaster* reference genome (Release 6, Dm6) was performed using Subread package. RNA-seq data analysis was performed according to a module designed by COMBINE-Australia (Github: https://github.com/COMBINE-Australia/RNAseq-R) in RStudio. FeatureCounts function in RSubread package was utilised to summarise reads into counts table in which features mapped to multiple genes were excluded. edgeR and limma packages were employed to analyse count data. To filter for lowly expressed genes, counts-per-million (CPM) were calculated and genes with CPM<2 (equivalent to count<10) were discarded. To remove composition biases between libraries, TMM normalisation was applied. To realise differential expression, counts data was transformed into log_2_CPM using the voom function in the limma package. Individual gene expression is visualised as Normalised log2 expression i.e. log_2_CPM. The empirical Bayes method was used to perform empirical Bayes shrinkage on the variances, estimate moderated t-statistics and generate p-values and log_2_(FC). To control for multiple testing during exploratory analysis of our RNA-seq data, the testing relative to a threshold (treat) function from the limma package was utilised to recalculate the moderated t-statistics and p-values for a FC cut-off of 1.5 and p-value of 0.05. RNA-seq results were plotted in RStudio using gplots, ggplot2, and VennDiagram packages. Perplexity AI (https://www.perplexity.ai/) was used to assist with R script troubleshooting, taking error prompts as inputs.

Gene Ontology (GO) enrichment analyses were performed on gene symbols extracted from org.Dm.eg.db package, using the topGO package in RStudio. Overrepresented GO terms were identified through Fisher’s exact test with weight method on genes with p-value<0.05 after treat. To realise GO enrichment, differential expressed genes were compared with those of all known genes present in the annotation. The results were plotted in RStudio using the ggplot2 package.

### Statistical analyses

At least three animals per genotype were used for all experiments. Statistical analyses and plotting were performed in GraphPad Prism 9. In graphs, data = mean ± standard error of the mean (SEM). n = lobes or VNC unless specified. For comparisons between two conditions, p-values were calculated by non-parametric Mann-Whitney tests when data show non-normally distribution. For comparisons between more than two conditions, p-values were calculated by Kruskal-Wallis tests for non-normally distributed data. Dunn’s tests were used to correct for multiple comparisons following Kruskal-Wallis tests. For comparisons of Imp and Syp intensities between *lola^E76^* and WT cells in the same brain, Wilcoxon matched-pairs signed rank test were used. (*) p<0.05 (**) p< 0.005, (***) p< 0.001, (****) p < 0.0001, (ns) p>0.05.

## Acknowledgements

We thank Lei Zhang, Helena Richardson, Kieran Harvey, Ward Odenwald, Alex Gould, Cedric Maurange, Tzumin Lee, Nicholas Sokol, Claude Desplan, Paul M Macdonald, Leonie Quinn and Kuniaki Saito for the generous sharing of antibodies and fly stocks. We thank Qian Dong for critical reading of the manuscript; Abdul Jabbar Saiful Hilmi and Sofya Golenkina for assisting with the generation of *UAS-chinmo^RNAi^* construct; Katrina Mitchell for assisting with RNA-seq data analyses; Perplexity AI for R script troubleshooting. We thank the Bloomington *Drosophil*a Stock Centre (BDSC), Vienna *Drosophila* Resource Centre (VDRC), and Developmental Studies Hybridoma Bank (DSHB) for fly stocks and antibodies. We thank OZDros for *Drosophila* quarantine. We thank Centre for Advanced Histology and Microscopy (RRID:SCR_025432), Research Flow Core (RRID:SCR_025550), and Molecular Genomics Core (RRID:SCR_025695) at Peter MacCallum Cancer Centre for access and support. We thank Australian *Drosophila* Transgenic Facility for injection service.

## Author contributions

PKN: Writing - original draft, Conceptualization, Investigation, Writing - review & editing, Methodology, Resources, Data curation, Validation, Formal analysis, Software, Visualization.

FF: Conceptualization, Methodology, Validation.

OM: Methodology.

TS: Methodology.

LYC: Writing-original draft, Conceptualization, Investigation, Writing - review & editing, Methodology, Resources, Funding acquisition, Data curation, Validation, Supervision, Formal analysis, Project administration, Visualization.

## Data availability

All data needed to evaluate the conclusions in the paper are present in the paper. The *UAS-chinmo^RNAi^*(II) *Drosophila* stock can be provided by Louise Y Cheng. Requests for this *Drosophila* stock should be submitted to. RNA-seq produced in this study is available at GEO (accession number: GSE306602).

