## Supplementary Figures for "Chinmo defines the region-specific oncogenic competence in the *Drosophila* central nervous system"

### Supplementary information

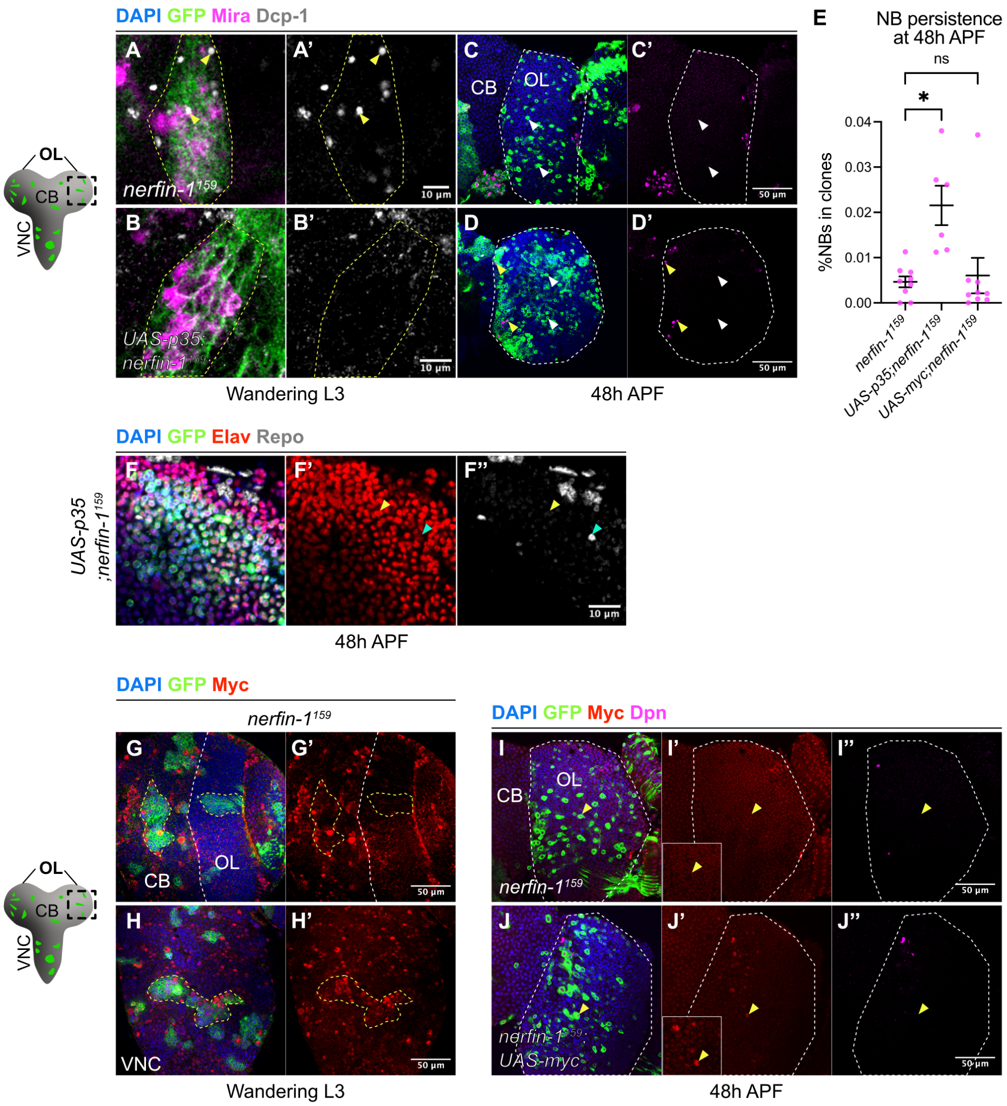


**Fig S1: The roles of apoptosis and Myc in the elimination of *nerfin-1*$^{159}$ ectopic NB**

**A-B'.** Single confocal sections of **(A-A')** *nerfin-1*$^{159}$ and **(B-B')** *UAS-p35;nerfin-1*$^{159}$ clones (dash lines) in the deep layers of the OLs at wandering L3 stage. DAPI (blue), GFP (green), Mira (magenta). Dcp-1 marks for apoptotic cells (grey, yellow arrowheads). Scale bars: 10 µm.

**C-D'.** Maximum projections of the OLs (dash lines) at 48h APF with **(C-C')** *nerfin-1*$^{159}$ and **(D-D')** *UAS-p35;nerfin-1*$^{159}$ clones. *UAS-p35* misexpression results in persisting *nerfin-1*$^{159}$ ectopic NBs (yellow arrowheads). White arrowheads indicate mutant cells devoid of Dpn. DAPI (blue), GFP (green), Dpn (magenta). Scale bars: 50 µm.

**E.** Quantifications of NB percentages in *nerfin-1*$^{159}$, *UAS-p35;nerfin-1*$^{159}$, and *UAS-myc;nerfin-1*$^{159}$ clones in the OLs at 48h APF. n=9, 6 and 9

**F.** Single confocal section of the OL at 48h APF with *UAS-p35;nerfin-1*$^{159}$ clones. DAPI (blue), GFP (green), Elav (red), Repo (grey). Yellow arrowhead indicates Elav$^+$Repo$^-$ mutant cell whereas cyan arrowhead indicates Elav$^-$Repo$^+$ mutant cell. Scale bars: 10 µm.

**G-H'.** Single confocal sections in the deep layers of **(F-F')** the brain lobe and **(G-G')** the VNC, with *nerfin-1*$^{159}$ clones (yellow dash lines) at wandering L3 stage. Myc (red) is expressed in *nerfin-1*$^{159}$ clones in the CB/VNC but not in the OLs. DAPI (blue), GFP (green). White dash lines indicate the CB/OL borders. Scale bars: 50 µm.

**I-J".** Maximum projections of the OLs (dash lines) at 48h APF with **(H-H")** *nerfin-1*$^{159}$ and **(I-I")** *UAS-myc;nerfin-1*$^{159}$ clones. DAPI (blue), GFP (green), Myc (red), Dpn (magenta). Scale bars: 50 µm.

Data information: Data are represented as mean ± SEM. *p*-values were obtained using Dunn's test to correct for multiple comparisons. $^*p < 0.05$.

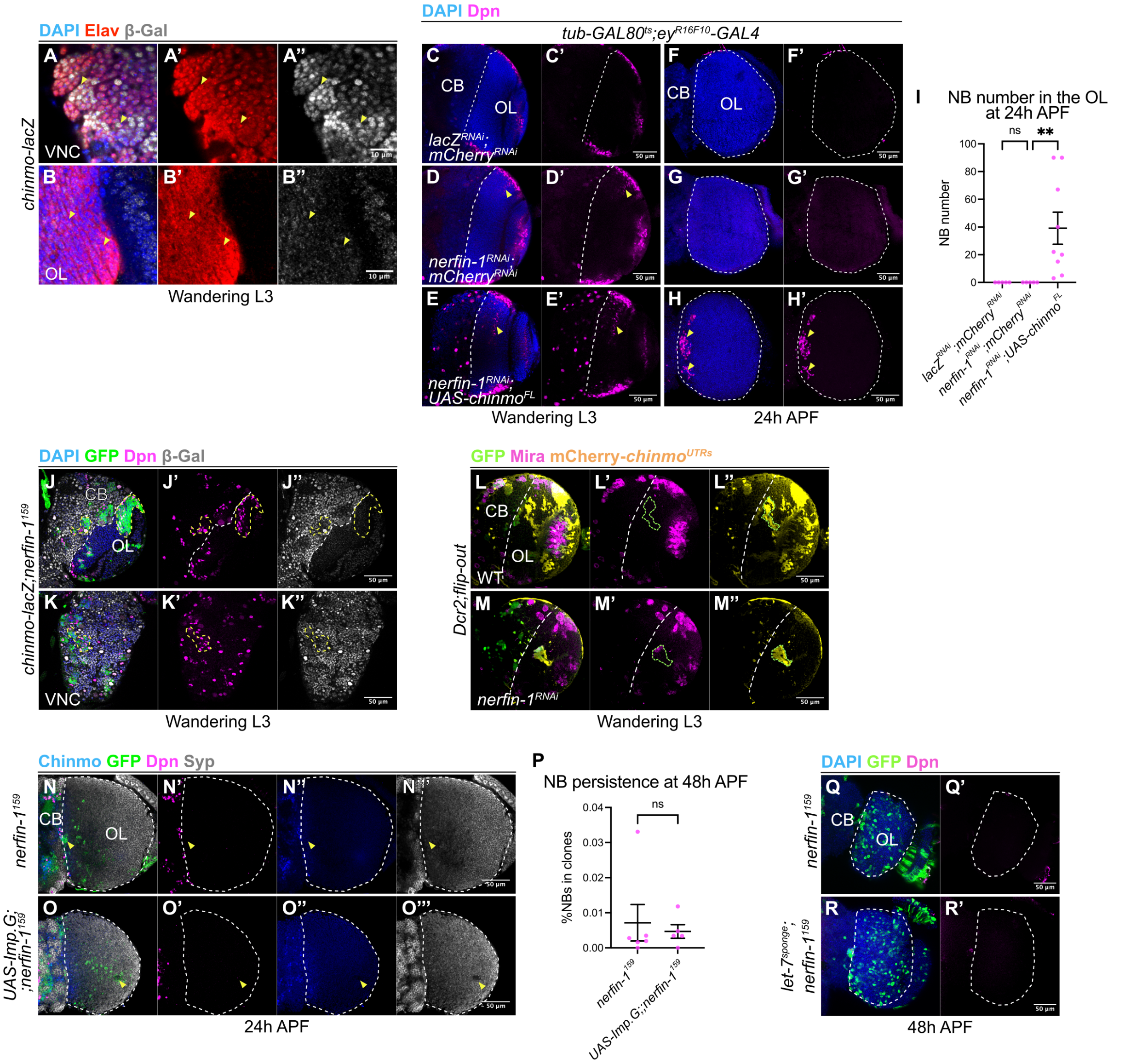


**Fig S2: Chinmo is not regulated at the post-transcriptional level in *nerfin-1$^{RNAi}$* clones in the OL; *nerfin-1$^{159}$* clones express EcR**

**A-B''.** Single confocal sections in the deep layers of **(A-A'')** VNC and **(B-B'')** OL expressing *chinmo*-lacZ at wandering L3 stage. DAPI (blue), Elav (red), β-Galactosidase (β-Gal, grey). Arrowheads indicate Elav$^{+}$ neurons. Scale bars: 10 µm.

**C-E'.** Single confocal sections in the deep layers of the brain lobes at wandering L3 stage with *tub-GAL80$^{ts}$;ey$^{R16F10}$-GAL4* driving **(C-C')** *lacZ$^{RNAi}$;mCherry$^{RNAi}$*, **(D-D')** *nerfin-1$^{RNAi}$;mCherry$^{RNAi}$*,and **(E-E')** *nerfin-1$^{RNAi}$;UAS-chinmo$^{FL}$*. DAPI (blue), Dpn (magenta). Arrowheads indicate ectopic NBs in the deep layers of the OL medulla. Dash lines indicate the CB/OL border. Scale bars: 50 µm.

**F-H'.** Maximum projections of the OLs (dash lines) at 24h APF with *tub-GAL80$^{ts}$;ey$^{R16F10}$-GAL4* driving **(F)** *lacZ$^{RNAi}$;mCherry$^{RNAi}$*, **(F-F')** *nerfin-1$^{RNAi}$;mCherry$^{RNAi}$*,and **(H-H')** *nerfin-1$^{RNAi}$;UAS-chinmo$^{FL}$*. DAPI (blue), Dpn (magenta). Arrowheads indicate ectopic NBs. Scale bars: 50 µm.

**I.** Quantification of NB numbers in the OL at 24h APF with *tub-GAL80$^{ts}$;ey$^{R16F10}$-GAL4* driving *lacZ$^{RNAi}$;mCherry$^{RNAi}$*, *nerfin-1$^{RNAi}$;mCherry$^{RNAi}$*,and *nerfin-1$^{RNAi}$;UAS-chinmo$^{FL}$*. n=5, 5 and 9.

**J-K''.** Single confocal sections in **(J-J'')** the brain lobe and **(K-K'')** the VNC at wandering L3 stage, with *chinmo-lacZ;nerfin-1$^{159}$* clones (yellow dash lines). White dash lines indicate the CB/OL border. DAPI (blue), GFP (green), Dpn (magenta), β-Gal (grey). Scale bars: 50 μm.

**L-M''.** Single confocal sections in the deep layers of the brain lobes with **(L-L'')** WT and **(M-M'')** *nerfin-1$^{RNAi}$* flip-out clones (green dash lines) at wandering L3 stage, both expressing *UAS-mCherry-chinmo$^{UTRs}$* (yellow), GFP (green), Mira (magenta). White dash lines indicate the CB/OL borders. Scale bars: 50 μm.

**N-O'''.** Single confocal sections of the OLs (dash lines) at 24h APF with **(N-N''')** *nerfin-1$^{159}$* and **(O-O''')** *UAS-Imp.G;;nerfin-1$^{159}$* clones. Chinmo (blue), GFP (green), Dpn (magenta), Syp (grey). Arrowheads indicate mutant cells. Scale bars: 50 μm.

**P.** Quantification of NB percentages in *nerfin-1$^{159}$* and *UAS-Imp.G;;nerfin-1$^{159}$* clones in the OLs at 48h APF. n=6 and 5.

**Q-R'.** Maximum projections of the OLs (dash lines) at 48h APF with **(Q-Q')** *nerfin-1$^{159}$* and **(R-R')** *let-7$^{sponge}$;nerfin-1$^{159}$* clones. DAPI (blue), GFP (green), Dpn (magenta). Scale bars: 50 μm.

Data information: Data are represented as mean ± SEM. *p*-values were obtained using Mann-Whitney test and Kruskal-Wallis test with Dunn's test to correct for multiple comparisons. $^{**}p < 0.005$.

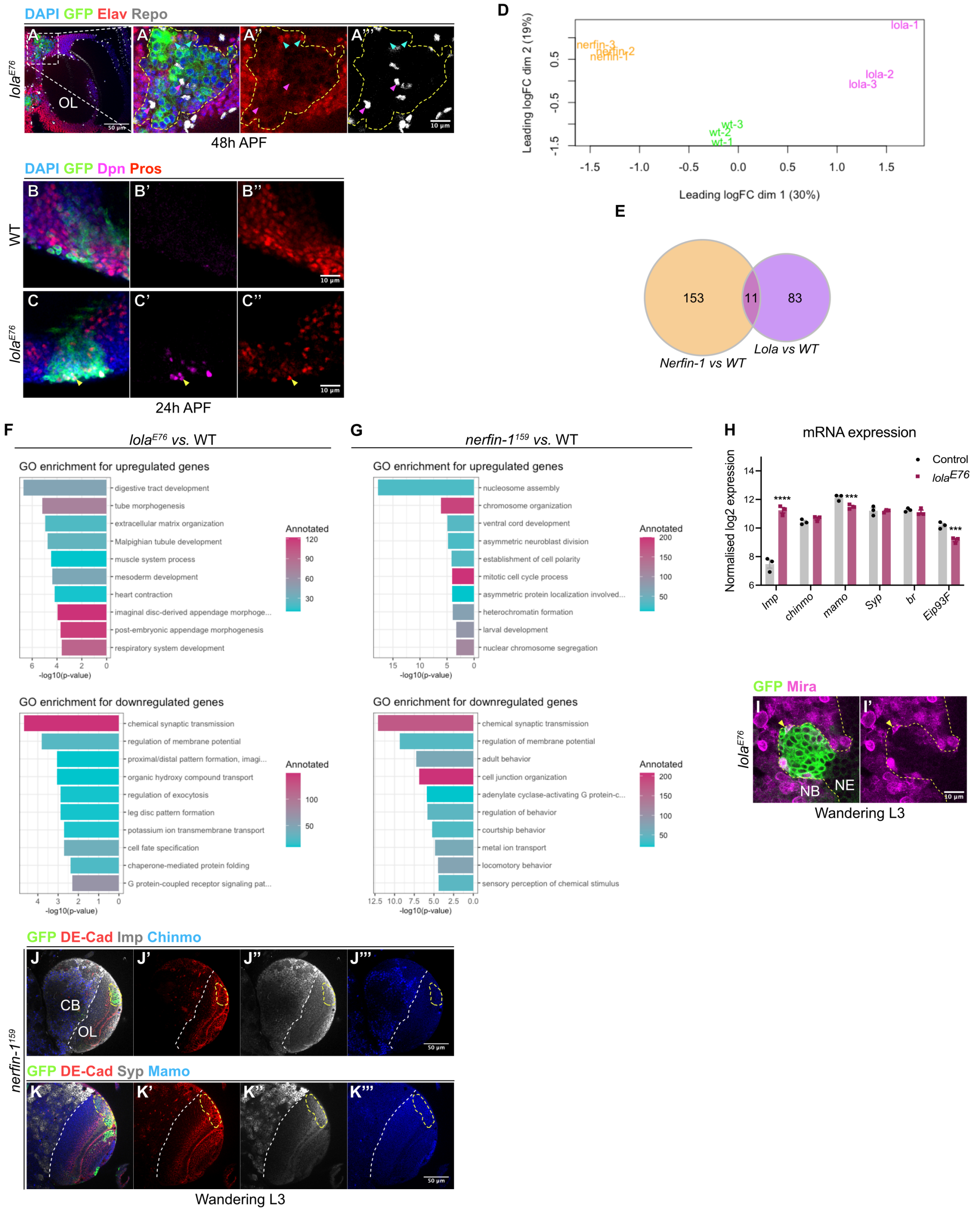


**Fig S3: RNA-seq identified minimal overlap in differentially expressed genes between Nerfin-1 and Lola; Nerfin-1 is dispensable for temporal progression in the larval medulla NE**

**A-A'''. (A)** Single confocal section of the OL at 48h APF with *lola*$^{E76}$ clones. Scale bars: 50 µm. **(A'-A''')** Boxed area in (A) Scale bars: 10 µm. Dash lines indicate mutant clone. DAPI (blue), GFP (green), Elav (red), Repo (grey). Cyan and magenta arrowheads indicate Elav$^{+}$GFP$^{+}$ and Repo$^{+}$GFP$^{+}$ cells, respectively.

**B-C''.** Single confocal section of **(B-B'')** WT and **(C-C'')** *lola*$^{E76}$ clones in the OL at 24h APF. Arrowheads indicate Dpn$^{+}$Pros$^{+}$ cells. DAPI (blue), GFP (green), Dpn (magenta), Pros (red). Scale bars: 10 µm.

**D.** Multidimensional plot of RNA-seq data of WT, *lola*$^{E76}$, and *nerfin-1*$^{159}$ clones collected from the late larval OLs.

**E.** Venn diagram indicating the degree of overlap in differentially expressed genes between *nerfin-1*$^{159}$ vs. WT, and *lola*$^{E76}$ vs. WT comparisons.

**F-G.** GO enrichment analyses of differentially expressed genes between **(F)** *lola*$^{E76}$ vs. WT cells and **(G)** *nerfin-1*$^{159}$ vs. WT cells from the late larval OLs.

**H.** Normalised log2 expression of the NE temporal factors *Imp, chinmo, mamo, Syp, br, Eip93F* from RNA-seq data of *lola*$^{E76}$ vs. WT clones from the late larval OLs. n=3, 3 and 3 samples.

**I-I'.** Single confocal section of *lola*$^{E76}$ clones (yellow arrowhead) in the superficial layers of the OL at wandering L3 stage. GFP (green), Mira (magenta). Dashed lines mark the border between the Mira$^-$ NE and Mira$^+$ NBs. Scale bars: 10 µm.

**J-K'''.** Single confocal sections in the deep layers of the brain lobes at wandering L3 stage containing *nerfin-1*$^{159}$ clones (yellow dash lines). GFP (green), DE-Cad (red). **(J-J''')** In *nerfin-1*$^{159}$ NE cells, Imp (grey) and Chinmo (blue) expression appear unaltered compared to WT surrounding cells. **(K-K''')** In *nerfin-1*$^{159}$ NE cells, the expression of Syp (grey) and Mamo (blue) appear unaltered compared to surrounding WT cells. White dash lines indicate the CB/OL borders. Scale bars: 50 µm.

Data information: Data are represented as mean ± SEM. *p*-values were obtained using moderated t-test. ****$p < 0.0001$, ***$p < 0.001$, **$p < 0.005$, *$p < 0.05$.

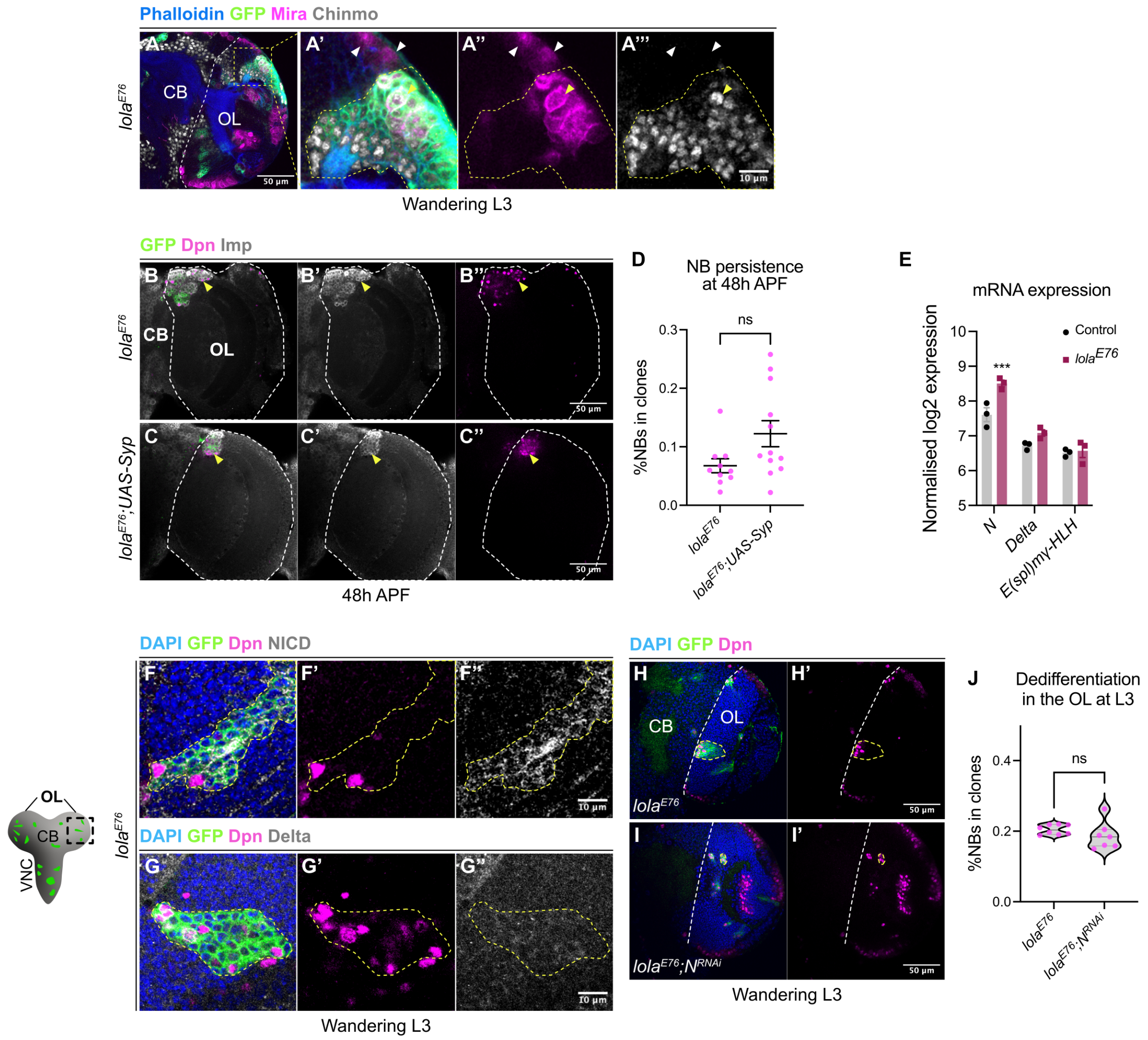


**Fig S4: Syp is not sufficient to inhibit Imp expression in *lola*$^{E76}$ clones; Notch signalling is not required for neuronal dedifferentiation induced by *lola* loss-of-function**

**A-A'''. (A)** Single confocal section in the deep layers of the brain lobe with *lola*$^{E76}$ clones (yellow dash lines). Scale bars: 50 µm. **(A'-A''')** Boxed area in (A). Scale bars: 10 µm. Phalloidin (blue), GFP (green), Mira (magenta), Chinmo (grey). White dash lines indicate the CB/OL border. White arrowheads indicate WT NBs, whereas yellow arrowheads indicate *lola*$^{E76}$ NBs.

**B-C''.** Maximum projections of the OLs (dash lines) at 48h APF with **(B-B'')** *lola*$^{E76}$ and **(C-C'')** *lola*$^{E76}$*;UAS-Syp* clones. Arrowheads indicate ectopic NBs. GFP (green), Dpn (magenta), Imp (grey). Scale bars: 50 µm.

**D.** Quantifications of NB percentages in *lola*$^{E76}$, and *lola*$^{E76}$*;UAS-Syp* clones in the OLs at 48h APF. n=10 and 12.

**E.** Normalised log2 expression of *N, Delta, E(spl)m*$\gamma$*-HLH* from RNA-seq data of *lola*$^{E76}$ vs. WT clones from the late larval OLs. n=3 and 3 samples.

**F-G''.** Single confocal sections of *lola*$^{E76}$ clones (dash lines) in the deep layers of the OL at wandering L3 stage, **(F-F'')** NICD (grey) and **(G-G'')** Delta (grey). DAPI (blue), GFP (green), Dpn (magenta). Scale bars: 10 µm.

**H-I'.** Single confocal sections in the deep layers of the brain lobes at wandering L3 stage with **(H-H')** *lola*$^{E76}$ and **(I-I')** *lola*$^{E76}$;*N*$^{RNAi}$ clones (yellow dash lines). DAPI (blue), GFP (green), Dpn (magenta). White dash lines indicate the CB/OL borders. Scale bars: 50 μm.

**J.** Quantifications of NB percentages in *lola*$^{E76}$ and *lola*$^{E76}$;*N*$^{RNAi}$ clones in the medulla at wandering L3 stage. n=8 and 7 clones.

Data information: Data are represented as mean ± SEM. *p*-values were obtained using Mann-Whitney test, moderated t-test. ***$p < 0.001$.

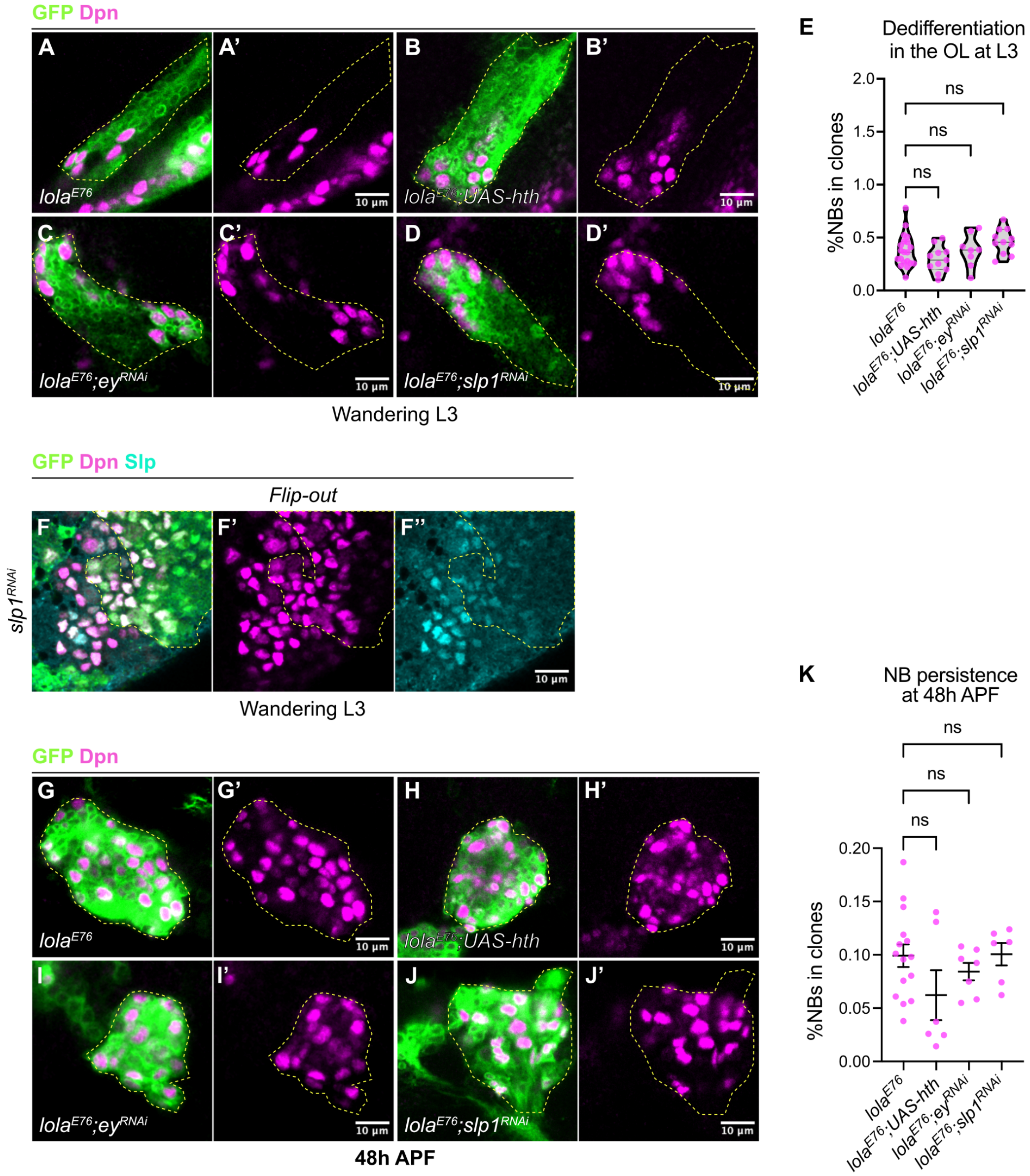


**Fig S5: The expression of NB tTFs is dispensable for the persistence of *lola*$^{E76}$ ectopic NBs in the pupal OL**

**A-D'.** Single confocal sections of **(A-A')** *lola*$^{E76}$, **(B-B')** *lola*$^{E76}$*;UAS-hth,* **(C-C')** *lola*$^{E76}$*;ey*$^{RNAi}$, and **(D-D')** *lola*$^{E76}$*;slp1*$^{RNAi}$ clones (dash lines) in the deep layers of the OL at wandering L3 stage. GFP (green), Dpn (magenta).

**E.** Quantification of NB percentages in *lola*$^{E76}$, *lola*$^{E76}$*;UAS-hth,* *lola*$^{E76}$*;ey*$^{RNAi}$, and *lola*$^{E76}$*;slp1*$^{RNAi}$ clones in the OL at wandering L3 stage. n=20, 11, 8 and 11 clones.

**F-F''.** Single confocal section of *slp1*$^{RNAi}$ flip-out clones (dash lines) in the superficial layers of the OL at wandering L3 stage. GFP (green), Dpn (magenta), Slp (cyan).

**G-J'.** Single confocal sections of **(G-G')** *lola*$^{E76}$, **(H-H')** *lola*$^{E76}$*;UAS-hth,* **(I-I')** *lola*$^{E76}$*;ey*$^{RNAi}$, and **(J-J')** *lola*$^{E76}$*;slp1*$^{RNAi}$ clones (dash lines) in the OL at 48h APF. GFP (green), Dpn (magenta).

**K.** Quantification of NB percentages in *lola*$^{E76}$, *lola*$^{E76}$*;UAS-hth, lola*$^{E76}$*;ey*$^{RNAi}$, and *lola*$^{E76}$*;slp1*$^{RNAi}$ clones in the OL at 48h APF. n=15, 6, 7 and 6 clones.

Data information: Data are represented as mean ± SEM. *p*-values were obtained using Kruskal-Wallis test and Dunn's test to correct for multiple comparisons. Scale bars: 10 μm.

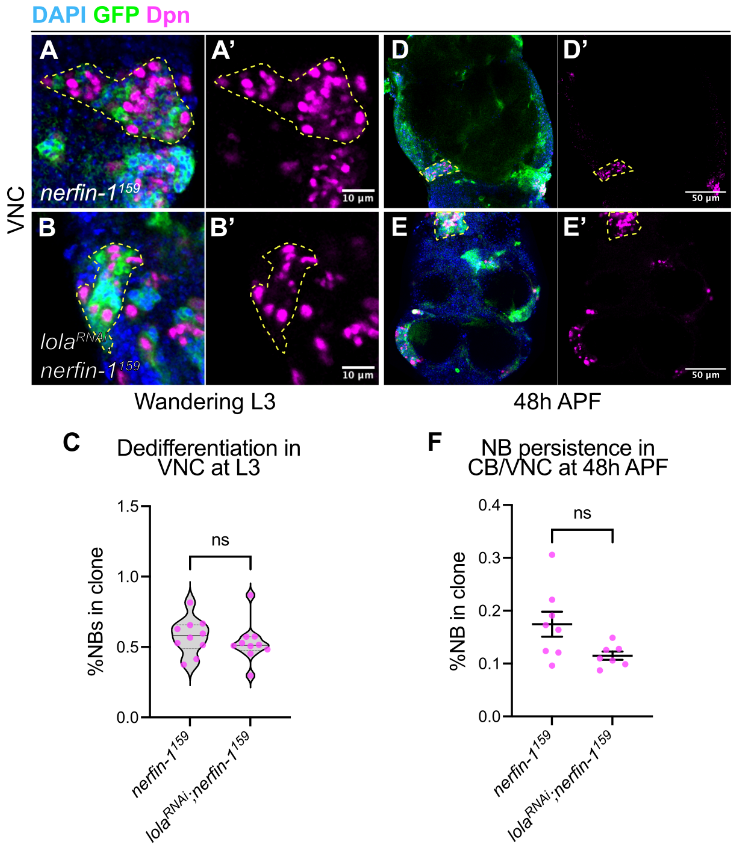


**Fig S6: Lola is not required for *nerfin-1*$^{159}$ ectopic NB persistence in the pupal VNC**

**A-B'.** Single confocal sections of **(A-A')** *nerfin-1*$^{159}$ and **(B-B')** *lola*$^{RNAi}$;*nerfin-1*$^{159}$ clones (dash lines) in the VNC at wandering L3 stage. DAPI (blue), GFP (green), Dpn (magenta). Scale bars: 10 µm.

**C.** Quantification of NB percentages in *nerfin-1*$^{159}$ and *lola*$^{RNAi}$;*nerfin-1*$^{159}$ clones in the VNC at wandering L3 stage. n=10 and 10 clones.

**D-E'.** Single confocal sections of the VNC at 48h APF with **(D-D')** *nerfin-1*$^{159}$ and **(E-E')** *lola*$^{RNAi}$;*nerfin-1*$^{159}$ clones (dash lines). DAPI (blue), GFP (green), Dpn (magenta). Scale bars: 50 µm.

**F.** Quantification of NB percentages in *nerfin-1*$^{159}$ and *lola*$^{RNAi}$;*nerfin-1*$^{159}$ clones in the VNC at 48h APF. n=8 and 7.

Data information: Data are represented as mean ± SEM. *p*-values were obtained using Mann-Whitney test.
